# ADAR2-mediated RNA editing of DNA:RNA hybrids is required for DNA double strand break repair

**DOI:** 10.1101/2021.03.24.436729

**Authors:** Sonia Jimeno, Rosario Prados-Carvajal, María Jesús Fernández-Ávila, Sonia Silva, Domenico Alessandro Silvestris, Martín Endara-Coll, Judit Domingo-Prim, Fernando Mejías-Navarro, Guillermo Rodríguez-Real, Amador Romero-Franco, Silvia Jimeno-González, Sonia Barroso, Valeriana Cesarini, Andrés Aguilera, Angela Gallo, Neus Visa, Pablo Huertas

## Abstract

The maintenance of genomic stability requires the coordination of multiple cellular tasks upon the appearance of DNA lesions. RNA editing, the post-transcriptional sequence alteration of RNA, has a profound effect on cell homeostasis, but its implication in the response to DNA damage was not previously explored. Here we show that, in response to DNA breaks, an overall change of the Adenosine-to-Inosine RNA editing is observed, a phenomenon we call the <u>R</u>NA <u>E</u>diting <u>DA</u>mage <u>R</u>esponse (REDAR). REDAR relies on the checkpoint kinase ATR and the recombination factor CtIP. Moreover, depletion of the RNA editing enzyme ADAR2 renders cells hypersensitive to genotoxic agents, increases genomic instability and hampers homologous recombination by impairing DNA resection. Such a role of ADAR2 in DNA repair goes beyond the recoding of specific transcripts, but depends on ADAR2 editing DNA:RNA hybrids to ease their dissolution.

**One sentence summary:** DNA recombination requires RNA editing of DNA:RNA hybrids to ease their melting and facilitate DNA end resection

## INTRODUCTION

Cells are continuously challenged by DNA damage. Among all kinds of insults that a DNA molecule has to deal with, double strand breaks (DSBs) are the most dangerous. Indeed, just one unrepaired DSB is enough to either kill or terminally arrest cells. For these reasons when DSBs are formed, a complex cellular response – the DNA damage response (DDR) - is triggered in order to ensure the proper repair of such a threat to genomic integrity^1^.

There are several pathways that can be used in order to repair a DSB and the choice between them is highly regulated. An eukaryotic cell can repair a DSB either by the simple re-ligation of the DNA ends (a process known as Non-Homologous End-Joining, NHEJ)^2^ or by a homology-driven repair event. There are different routes among the repair pathways that use homologous regions for repair, all of which are grouped in a process called homologous recombination (HR)^3^. All HR events share a first biochemical step called DNA resection, which is key to decide the pathway that will be eventually used to repair the DSB^4,5^. This process consists of the nucleolytic degradation of the DNA ends of the break that produces tails of 3’ended single stranded DNA (ssDNA) that are rapidly protected by the RPA protein complex.

In recent years, the importance of RNA and RNA-related factors in DNA repair has become clear^6–9^. Indeed, many RNA-related proteins have been shown to be targets of the DNA damage-induced post-translational modifications^10–12^. Also, direct roles of specific RNA-related factors in DNA repair have been recently reported (for a review see ^9^). Moreover, the RNA molecule itself seems to impact on DNA repair. Several labs have shown the formation of DNA:RNA hybrids around DSBs in different eukaryotes, either dependent on previous transcription^13,14^ or upon *de novo* transcription of the broken chromatin ^15,16^. The relevance of such RNA molecules is still under debate, with both pro- and anti-repair effects ascribed to them^9^.

An important co-transcriptional RNA modification that, so far, has not been extensively studied in its putative relationship with DNA repair and the response to DNA damage is RNA editing. This process alters RNA sequences by the action of specific deaminases that convert one base into another. Every mammalian transcript can be subjected to RNA editing^17–19^. RNA editing can be classified in several categories^20^, including Adenosine-to-Inosine (A-to-I) deamination, which is accomplished by a family of RNA-Specific Adenosine Deaminase known as ADARs^18,19^. This family is formed by ADAR1, ADAR2 (also known as ADARB1) and ADAR3; however, only ADAR1 and ADAR2 have been shown to present catalytic activity. A-to-I deamination is the most abundant form of RNA-editing in mammals and defects in this process are associated with human diseases, such as disorders of the central nervous system^21^ or paediatric astrocytomas^22^. Only limited information has been published regarding the connection of A-to-I editing and DNA damage, albeit at least the mRNA of NEIL1, has been shown to be re-coded by ADAR1 to alter its enzymatic properties^23^. Moreover, A-to-I editing has been proposed to be involved in the pathogenesis of cancer^24,25^.

Here we show that the general pattern of ADAR2-mediated A-to-I editing changes upon DSB formation. Such changes depend on the DDR, specifically the ATR kinase and the resection protein CtIP. As a consequence, ADAR2 is required for maintenance of genomic integrity and, specifically for DNA end resection and HR. Strikingly, mRNAs from either resection-related or recombination-related genes are not affected by ADAR2. Instead, ADAR2 role in resection is related with its ability to edit DNA:RNA hybrids. Not only such structures increase when ADAR2 is depleted, but this protein physically and functionally interacts with the BRCA1-SETX complex for this role.

## EXPERIMENTAL PROCEDURES

### Cell lines and growth conditions

All cell lines were grown in DMEM (Sigma-Aldrich) supplemented with 10% fetal bovine serum (Sigma-Aldrich), 2 mM L-glutamine (Sigma-Aldrich), 100 units ml^−1^ penicillin and 100μg ml^−1^ streptomycin (Sigma-Aldrich). U2OS and U118-derived cell lines stably expressing GFP or GFP-ADAR2 plasmids ^26^ were grown in standard medium supplemented with 0.5 mg ml^−1^ G418 (Gibco, Invitrogen). Cells expressing RNWG and RNAG were grown in in standard medium supplemented with 0.5 mg ml^−1^ G418 (Gibco, Invitrogen).

### siRNAs, plasmids and transfections

siRNA duplexes were obtained from Sigma-Aldrich or Dharmacon (Supplementary Table S2) and were transfected using RNAiMax Lipofectamine Reagent Mix (Life Technologies), according to the manufacturer’s instructions. RNWG and RNAG was a gift from Dr. Jantsch’s lab^27^. The GFP-ADAR2 and GFP-ADAR2 mutant (GFP-ADAR2-E/A-) plasmids were previously described^26^. RNaseH1 overexpression was achieved with the pCDNA3-RNAseH1 vector^28^, and pCDNA3 (Invitrogen) was used as a control. Plasmid transfection of U2OS cells was carried out using FuGENE 6 Transfection Reagent (Promega) according to the manufacturer’s protocol, with the exception of pCDNA3-RNaseH1 and pCDNA3 plasmids that were transfected using Lipofectamine 3000 (Invitrogen) according to the manufacturer’s instructions.

### HR and NHEJ analysis

U2OS cells bearing a single copy integration of the reporters DR-GFP (Gene conversion)^29^, SA-GFP (SSA)^30^ or EJ5-GFP (NHEJ)^30^ were used to analyse the different DSB repair pathways. In all cases, 50,000 cells were plated in 6-well plates in duplicate. One day after seeding, cells were transfected with the indicated siRNA and the medium was replaced with fresh one 24h later. The next day, each duplicate culture was infected with lentiviral particles containing I-SceI–BFP expression construct at MOI 10 using 8 μg/ml polybrene in 1.5 ml of DMEM. Then, cells were left to grow for an additional 24 h before changing the medium for fresh DMEM. 48h after siRNA transfection, cells were washed with PBS, trypsinised, neutralized with DMEM, centrifuged for 5 min at 700 g, fixed with 4% paraformaldehyde for 20 min and collected by centrifugation. Then, cell pellets were washed once with PBS before resuspension in 150 μl of PBS. Samples were analysed with a BD FACSAria with the BD FACSDiva Software v5.0.3. Four different parameters were considered: side scatter (SSC), forward scatter (FSC), blue fluorescence (407 nm violet laser BP, Filter 450/40), green fluorescence (488 nm blue laser BP Filter 530/30). Finally, the number of green cells from at least 10,000 events positives for blue fluorescence (infected with the I-SceI–BFP construct) was scored. The average of both duplicates was calculated for each sample of every experiment. To facilitate the comparison between experiments, this ratio was normalized with siRNA control. At least four completely independent experiments were carried out for each condition and the average and standard deviation is represented.

### RNA editing assay in *vivo*

Cells were seeded in 60 mm plates and transfected with siRNAs 24h later. Cells were irradiated with 10 Gy or treated with the indicated doses of CPT and incubated during 12h before harvesting. In the experiments performed with protein inhibitors, the medium was exchanged 2 h before irradiation with fresh DMEM containing 10 μM ATMi (KU55933), 5 μM ATRi (ETP46464)), 10 μM DNA-PKi (NU7441) or DMSO as control. After that, the medium was replaced, DSBs were induced with the indicated DNA damage agent and cells were incubated for 10 hours. Then, cells were harvested with trypsin, spun down at 500 g for 5 min and washed with PBS. Cells were fixed with 4% paraformaldehyde for 15 min at 4°C in the dark, and later rinsed and resuspended in 150 μl of PBS. Red and green fluorescence was measured on BD FACSAriaTM using FACSDiva v5.0.3 software as indicated in above section.

#### Clonogenic cell survival assays

To study cell survival after DNA damage, clonogenic assays were carried out seeding cells in 6-well plates at two different concentrations in triplicates. DSBs were produced by IR or by acute treatment with topoisomerase inhibitor camptothecin (CPT; Sigma). For IR, 250 and 500 transfected cells were seeded per well and, for drug treatments, 500 and 1,000 cells per well. The following day, cells were exposed to DNA damaging agents: 2 Gy, 4 Gy or mock treated or incubated for 1h with 0.01, 0.05 or 0.1 μM CPT or vehicle (DMSO) as control. After two washes with PBS, fresh medium was added and cells were incubated at 37°C for 7-14 days to allow colony formation. Afterwards, cells were stained and visualized in solution of 0.5% Crystal Violet (Merck) and 20% ethanol (Merck). Once the colonies were stained, this solution was removed and plates were washed with water. The surviving percentage at each dose was calculated by dividing the average number of visible colonies in treated versus control (untreated or vehicle-treated) dishes.

### SDS-PAGE and western blot analysis

Protein extracts were prepared in 2× Laemmli buffer (4% SDS, 20% glycerol, 125 mM Tris-HCl, pH 6.8) and passed 10 times through a 0.5 mm needle–mounted syringe to reduce viscosity. Proteins were resolved by SDS-PAGE and transferred to low fluorescence PVDF membranes (Immobilon-FL, Millipore). Membranes were blocked with Odyssey Blocking Buffer (LI-COR) and blotted with the appropriate primary antibody and infra-red dyed secondary antibodies (LI-COR) (Table S3 and S4). Antibodies were prepared in blocking buffer supplemented with 0.1% Tween-20. Membranes were air-dried in the dark and scanned in an Odyssey Infrared Imaging System (LI-COR), and images were analysed with ImageStudio software (LI-COR).

### Immunoprecipitation

U2OS cells or U2OS cells containing GFP or GFP-CtIP were harvested in lysis buffer (50 mM Tris-HCl, pH 7.4, 100 mM NaCl, 1 mM EDTA, 0.2 % Triton X-100, 1X protease inhibitors (Roche), 1X phosphatase inhibitor cocktail 1 (Sigma)) and incubated for 30 min on ice with Benzonase (90 U/ml), with the exception of the S9.6 IP in which samples were sonicated for 15 min. Protein extract (1 mg) was incubated at 4 °C with 10 μl of anti-BRCA1, anti-SETX, anti-ADAR2 or S9.6 antibody (Supplementary Table S3) or with an equivalent amount of IgG (Mouse of Rabbit) as negative control. Afterwards, extracts were incubated with magnetic protein A Dynabeads (Novex) overnight. Beads were then washed three times with lysis buffer, and the precipitate was eluted in 50 μl of Laemmli buffer 2x.

### Immunofluorescence and microscopy

For RPA, γH2AX and BRCA1 foci visualization, U2OS cells knocked-down for different proteins were seeded on coverslips. Cells were treated with 10 Gy ionizing irradiation and incubated for 1 h. Then, coverslips were washed once with PBS followed by treatment with Preextraction Buffer (25 mM Tris-HCl, pH 7.5, 50 mM NaCl, 1 mM EDTA, 3 mM MgCl_2_, 300 mM sucrose and 0.2% Triton X-100) for 5 min on ice. Cells were fixed with 4% paraformaldehyde (w/v) in PBS for 20 min. For 53BP1 foci, cells growing on coverslips were treated for 10 min on ice with methanol and 30 s with acetone. In all cases, following two washes with PBS, cells were blocked for 1 h with 5% FBS in PBS, co-stained with the appropriate primary antibodies (Supplementary Table S3) in blocking solution overnight at 4°C or for 2 h at room temperature, washed again with PBS and then co-immunostained with the appropriate secondary antibodies (Supplementary Table S4) in blocking buffer. After washing with PBS and dried with ethanol 70% and 100% washes, coverslips were mounted into glass slides using Vectashield mounting medium with DAPI (Vector Laboratories). Images were acquired and analysed using a Leica Fluorescence microscope. The analysis of 53BP1 number foci formation was performed automatically using MetaMorph software.

### SMART (Single-Molecule Analysis of Resection Tracks)

SMART was performed as described ^31^. Briefly, cells were grown in the presence of 10 μM BrdU for less than 24 h. Cultures were then irradiated (10 Gy) and harvested after 1 h. Cells were embedded in low-melting agarose (Bio-Rad), followed by DNA extraction. DNA fibres were stretched on silanized coverslips, and immunofluorescence was carried out to detect BrdU (Supplementary Table S3 and S4). Samples were observed under a Nikon NI-E microscope, and images were taken and processed with the NIS ELEMENTS Nikon Software. For each experiment, at least 200 DNA fibres were analysed, and the length of the fibers was measured with Adobe Photoshop CS4.

### Cell cycle analysis

Cells were fixed with cold 70% ethanol overnight, incubated with 250 μg ml^−1^ RNase A (Sigma) and 10 μg ml^−1^ propidium iodide (Fluka) at 37°C for 30 min and analysed with a FACSCalibur (BD). Cell cycle distribution data were further analysed using ModFit LT 3.0 software (Verity Software House Inc).

### UV laser micro-irradiation

Cells were micro-irradiated using a wide-field Angström’s microscope (Leica) equipped with a Micropoint pulsed dye laser of 365 nm (Photonic instruments, Inc.). The cells were seeded in 25mm coverslips and cultured overnight in the presence of 10 μM BrdU before laser micro-irradiation. About 40-50 cells were micro-irradiated with one laser stripe per cell. For RPA study, cells were pre-permeabilized with CSK Buffer (10 mM PIPES, 300 mM sucrose, 100 mM NaCl, 3 mM MgCl2, 1 mM EGTA) for 10 min. Then, cells were fixed for 10 min with 3.6% formaldehyde, permeabilized with 0.1% Triton X-100 for 15 min, and blocked for 30 min with 5% bovine serum albumin (BSA) in phosphate buffered saline (PBS). Antibodies (Supplementary Tables S3 and S4) were diluted in 1% BSA in PBST (PBS containing 0.01% Tween-20) and incubated for 1 h. Coverslips were mounted using Vectashield mounting medium with DAPI (VectorLabs) and the slides were visualized in a LSM780 confocal microscope (Carl Zeiss) with an optical thickness of 0.9 μm. Quantitative analyses of the number of cells with foci or stripes were carried out in random areas using FIJI software. The number of stripes was quantified in 20–30 cells per preparation.

### RNA isolation, RNA sequencing and *in silico* analysis

RNA extracts were obtained from cells using RNasey Mini kit (QIAGEN, 74104) according to manufacturer’s instructions. The RNA was purified with a standard phenol:chloroform extraction followed by an ethanol precipitation.

RNA concentration was quantified by measuring 260 nm absorbance using a NanoDrop ND-1000 spectrophotometer, and the quality of the sample was checked by running a 1% agarose gel and by RNA 6000 Nano assay on a 2100 Bioanalyzer (Agilent Technologies).

RNA-Seq data (76 bp strand-oriented reads generated from Illumina platform) were first processed for adaptors trimming and low-quality reads filtering. Then, cleaned reads were mapped against reference human genome (hg19), transcriptome and dpSNP by HISAT2 v.2.0.1 ^32^ and only uniquely and concordantly mapped reads have been used for subsequent analyses. RNA editing analysis was performed using a specific Python tool, REDItools^33^, with default parameters for the detection of the RNA editing sites collected in REDIportal database^34^. Recoding Editing Index was calculated as previously shown^35^.

### DNA:RNA hybrid detection

S9.6 (hybridoma cell line HB-8730) and RNase H1 (15606-1-AP; Proteintech) immunofluorescence (IF) was performed 48 h after siRNA transfection. Cells were fixed with ice-cold methanol, blocked and subsequently incubated with the primary and secondary antibodies. Nuclei were stained with DAPI. The S9.6 signal in nucleoli was subtracted from the integrated nuclear S9.6 signal to perform the analysis. Immunofluorescence images were acquired using a Leica DM6000 wide-field microscope equipped with a DFC390 camera at x63 magnification using the LAS AF software (Leica). Microscopy data analysis was performed using the Metamorph v7.5.1.0 software (Molecular Probes).

### ICGC data retrieval and analysis

Mutations sets were retrieved from the International Cancer Genome Consortium (ICGC) Data Portal MALY-DE and CLLE-ES datasets. ADAR2 expression levels from each donor were obtained using the UCSC Xena web tool. Open-access somatic mutations information from each mutation set was obtained by comparing each mutation set with the latest release of the Aggregated Somatic Mutation VCF file by the ICGC using custom python scripts. The percentage of A to G and T to C mutations was calculated as the quotient between the number of A to G mutations and T to C mutations to the total number of mutations from each donor set.

### Statistical analysis

Unless specifically specified, statistical significance was determined with a Student’s t-test using PRISM software (Graphpad Software Inc.). Statistically significant differences were labelled with one, two or three asterisks if *p* < 0.05, *p* < 0.01 or *p* < 0.001, respectively.

## RESULTS

### RNA editing changes after DNA damage

As previously mentioned, a crosstalk between RNA metabolism and DNA repair has been extensively documented^9^, but a connection between DNA repair and RNA editing has not been extensively analysed. Thus, we wanted to study whether the appearance of DNA damage had any effect on RNA editing. In order to explore this possibility, we used a previously published reporter system (RNAG) that measures levels of RNA editing using the accumulation of the fluorescent proteins GFP and RFP^27^. This system bears both the RFP and GFP ORFs in a single transcript, with a stop codon between them (Figure 1A). So, cells bearing such reporter express RFP constitutively, but GFP is only produced if an RNA editing event changes the A of the stop codon to an I (Figure 1A)^27^. Therefore, the number of red cells that are also green indicates the efficiency of RNA editing. As a control to discard other effects non-related to editing on this system, we used the RNWG control reporter, in which the stop codon is pre-edited so all cells bearing the construct fluoresce, indeed, in red and green^27^. In U2OS cells stably transfected with the reporter we observed that DNA damage induced by ionizing radiation increased GFP expression by 50% specifically in the RNAG reporter and not in the RNWG control, in agreement with a DNA damage-stimulation of RNA A-to-I editing in this system (Figure 1B). Similar results were obtained when using the DNA damage inducing drug camptothecin (Figure 1C), where we could observe a dose-dependent effect on RNA editing stimulation. One possibility is that DNA damage induces the accumulation of the A-to-I editing machinery, namely ADAR1 and ADAR2 enzymes, thus increasing this process. However, neither of these proteins was upregulated, but slightly downregulated, upon exposure to IR (Supplementary Figure S1A). Then, in order to confirm this was a canonical induction of RNA editing, we depleted the A-to-I editing machinery. To choose which member of the ADAR family to downregulate, we revisited the data we obtained in a previous genome-wide screening for factors that unbalance the choice between DSB repair pathways^36^. Interestingly, both ADAR1 and ADAR2, but not the catalytically inactive ADAR3, skewed DSB repair towards end-joining (Supplementary Figure S1B), and this was not due to changes in the cell cycle (Supplementary Figure S1C). However, the effect was more prominent and clearer upon ADAR2 depletion. Indeed, downregulation of ADAR2 severely compromised both the basal and the DNA damage-induced expression of the GFP in the RNAG (Figure 1D; for ADAR2 depletion efficiency see Supplementary Figure S2A), but, as expected, not in the RNWG control reporter (Supplementary Figure S2B).

**Figure 1.**
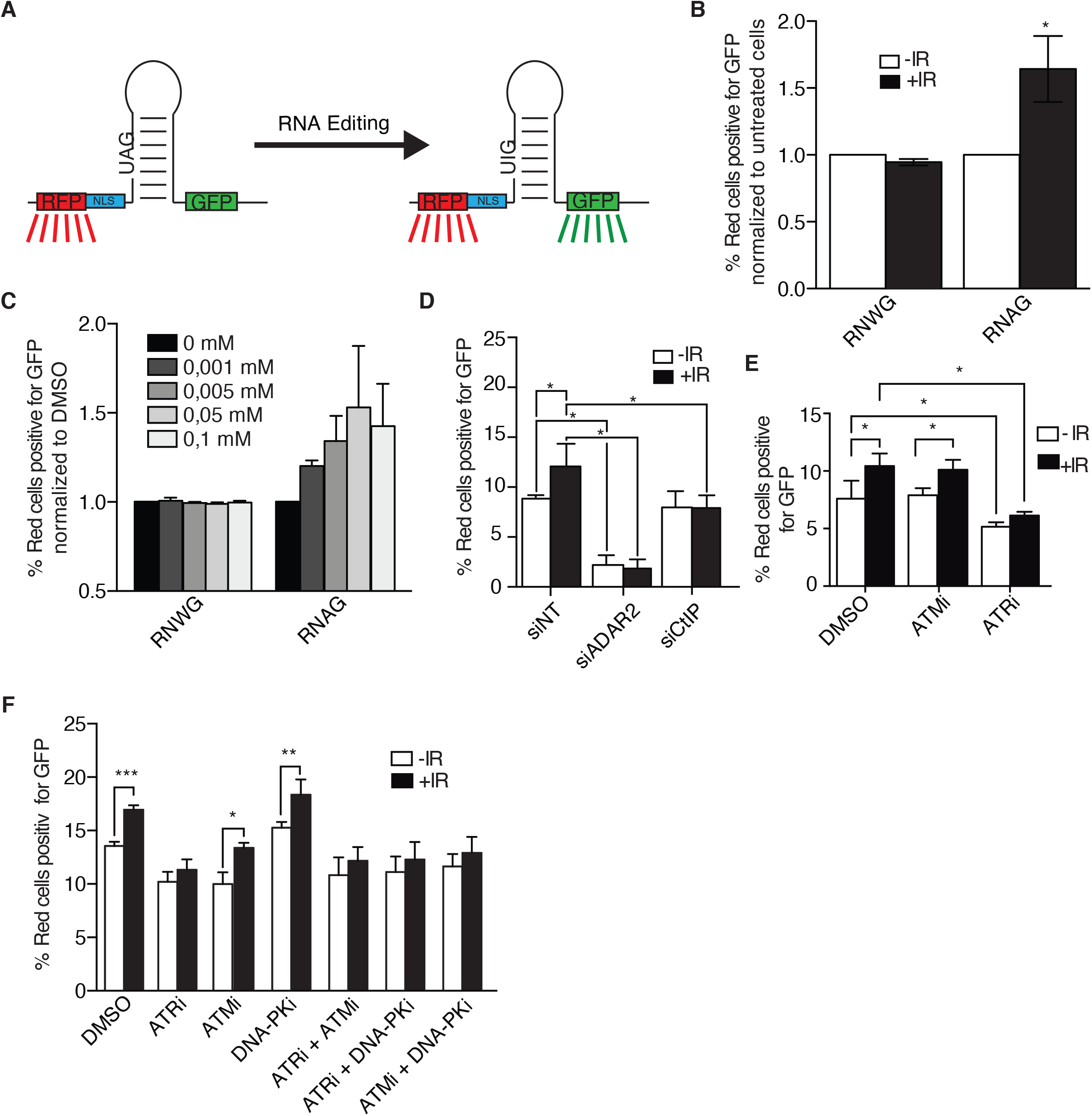
DNA damage increases RNA editing. (**A)** Scheme of the RNAG editing system. A bi-cistronic mRNA containing the RFP and GFP sequences is produced. The presence of a stop codon impedes the expression of the GFP ORF, except when the adenine is edited to inosine. The presence of a secondary structure containing such stop codon allows its recognition and deamination of the adenine by ADAR proteins. **(B**) DNA damage induced RNA editing. The plot shows the percentage of cells bearing the RNAG reporter or the constitutively edited RNWG system that express both the RFP (red cells) and GFP (green cells). Cells were either irradiated (10 Gy; black bars) or mock treated (white bars) and incubated for 12 h. The percentage of green cells over 10.000 red cells were analysed on BD FACSAriaTM using FACSDiva v5.0.3 software. For each reporter, the ratio of green and red cells was normalized with the untreated conditions. (**C)** Same as **B**, but cells were treated with the indicated concentration of CPT. **(D)**. Cells bearing the RNAG reporter were transfected with the indicated siRNAs and irradiated or not, and the percentage of red cells that were also green is plotted. (**E**) Same as **D**, but, cells were pretreated for 2 hours with 10 μM of inhibitors of ATM (ATMi), ATR (ATRi) or DMSO as control, previous to the irradiation. Cells were collected to check for editing levels 10 hours after irradiation. The inhibitors were kept for the duration of the experiment. (**F**) Same as **E**, but, cells were also treated with DNA-PK inhibitor (DNA-PKi), as well as the double combinations of the ATM, ATR and DNA-PK inhibitors, as indicated. The average and the standard deviation of the medians of at least four independent experiments are shown. Statistical significance was determined with a Student’s *t*-test. One, two or three asterisks represent p<0.05, p<0.01 or p<0.001, respectively.

To better understand this phenomenon, we decided to look for DDR factors that affected the DNA damage-induced RNA editing. Recently, we have found that CtIP, a core DNA end resection factor that is also required for ATR activation, plays additional roles in DNA damage-induced RNA splicing^37^. Interestingly, we could see that CtIP downregulation specifically eliminated the DNA damage-dependent induction of RNA editing without affecting the basal levels (Figure 1D). Again, CtIP depletion did not alter GFP levels in the control RNWG system (Supplementary Figure S2B). We could also complement this effect with the expression of siRNA-resistant flag-tagged CtIP in CtIP depleted cells, to the same extent as the control cells transfected with a non-targeting siRNA, even though overexpression of FLAG-CtIP on its own reduced the intensity of this DNA damage-induced phenotype (Supplementary Figure S2C).

The general response to DNA damage is mainly controlled by the activation of two related apical kinases, ATM and ATR^1^. Thus, we also tested if any of them was required for the induction of RNA editing upon irradiation. Interestingly, ATM inhibition did not affect DNA damage-induced RNA editing, while ATR inhibition decreased the DNA damage-induced editing increase (Figure 1E). This agrees with the notion that ATR and CtIP act on the same branch of the DNA damage checkpoint in response to DSBs^38^. The lack of response with the ATM inhibitor could be explained by a compensation by another member of the PIKK family, most likely DNA-PK. Along those lines, the ATR effect could also be affected by this phenomenon. Thus, we repeated the experiment with ATM, ATR and DNA-PK inhibitors in different combinations (Figure 1F). As shown, inhibition of ATR suppressed the DNA damage-induction of RNA editing, regardless of the presence of the inhibitors of ATM or DNA-PK. Interestingly, chemical inhibition of DNA-PK showed a limited increase in the basal levels of RNA editing, but importantly the exposure to DNA damage still provoked a hyper-activation of the process. Notably, concomitant inhibition of both DNA-PK and ATM abolished the induction of RNA editing caused by irradiation. Thus, it seems that those two kinases could have an overlapping role in this phenomenon.

### ADAR2 depletion causes genomic instability and DNA damage sensitivity

To understand the consequences of reduced A-to-I RNA editing for genomic stability, we decided to globally reduce such RNA modifications. Based on our previous data with ADAR2 (Figure 1D and Supplementary Figure S1B), we decided to use the downregulation of this protein as a tool to reduce A-to-I RNA editing. Strikingly, and in agreement with a role in maintaining genomic stability, the depletion of ADAR2 impaired DSB repair, measured as the presence of γH2AX foci 24h after irradiation. Spontaneous DNA damage accumulated in the absence of any exogenous genotoxic agent in ADAR2-depleted cells (Figure 2A). Confirming a DSB repair impairment, the disappearance of γH2AX foci upon exposure to ionizing radiation was delayed (Figure 2B). Indeed, repair levels of DSBs at 6 and 24 hours after irradiation in ADAR2-depleted cells were similar to those observed after downregulation of the critical repair factor CtIP (Figure 2B). Interestingly, and in agreement with an increased burden of spontaneous DNA damage, in the absence of ADAR2 we observed a significative increase of BRCA1 foci in cells unchallenged with any genotoxic agents (Figure 2C). A similar effect was observed upon ADAR1 downregulation (Figure 2D). Furthermore, micronuclei accumulated at high levels in ADAR2-depleted cells, regardless of the exposure to an external source of DNA damage (Figure 2E). Finally, and confirming a role of ADAR2 in DNA repair and the maintenance of genomic stability, its depletion rendered cells hypersensitive to DSBs-inducing agents such as ionizing radiation or camptothecin (Figures 2F and 2G).

**Figure 2.**
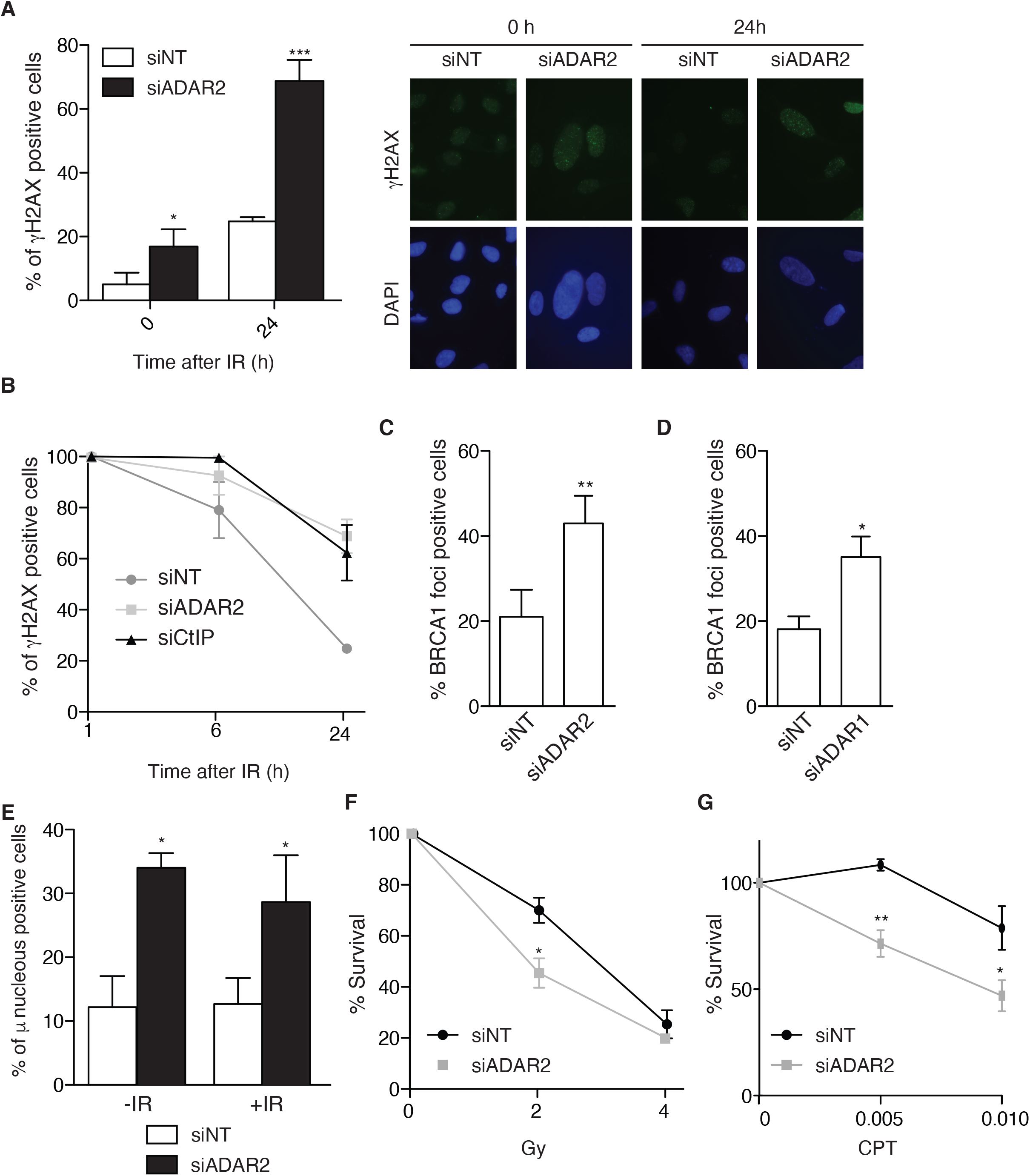
ADAR2 depletion causes genetic instability and DNA repair defects. **(A)** Percentage of cells positive for γH2AX foci upon spontaneous accumulation (0 h) or 24 hours post-irradiation with 10 Gy in U2OS cells transfected either a siRNA against ADAR2 or with control siNT. Quantification is shown on the left. A representative image is shown on the right. **(B)** Repair kinetics is shown as the disappearance of γH2AX foci 1-, 6- and 24-hours post-irradiation with 10 Gy in U2OS cells transfected either a siRNA against ADAR2, CtIP or with control siNT. **(C)** Percentage of spontaneous BRCA1 foci–positive cells in cells transfected with either a siRNA against ADAR2 or with control siNT. The average and SD of three independent experiments is shown. **(D)** Same as **C** but in cells depleted for ADAR1. **(E)** Percentage of cells positive for micronucleus without exposure to DNA damage (-IR) or 24 hours post-irradiation with 10 Gy (+IR) in U2OS cells transfected either with a siRNA against ADAR2 or with control siNT. Other details as in (A). **(F)** Clonogenic assays of U2OS cells depleted with a siRNA against ADAR2 or with control siNT after treatment with different doses of IR. Other details as in (A). **(G)** Same as **F** but cells treated with CPT (μM; right). In all panels, the average and SD of three independent experiments are shown and statistical significance was measured with a Student’s *t*-test. One, two or three asterisks represent p<0.05, p<0.01 or p<0.001, respectively.

### ADAR2 depletion affects DNA repair pathway choice

Next, we decided to test a possible requirement of A-to-I RNA editing for DSB repair. As mentioned, ADAR2 depletion skewed the balance between HR and NHEJ towards the latter (see reference^36^; Supplementary Figure S1B), suggesting that recombination might be compromised. Indeed, both RAD51-dependent gene conversion (GC) and RAD51-independent Single Strand Annealing (SSA), two types of homology-dependent repair (HDR), were reduced in cells downregulated for ADAR2 (Figures 3A and B). In stark contrast, there was no impact on NHEJ efficiency (Figure 3C), arguing that ADAR2 was particularly required for homologous recombination (HR). Cell cycle is a major regulator of DSB repair pathway choice, as HR is limited in G1. However, the observed HR defect was not caused by an accumulation of G1 cells (Supplementary Figure S1C). Thus, we conclude that ADAR2 facilitates repair by homologous recombination.

**Figure 3.**
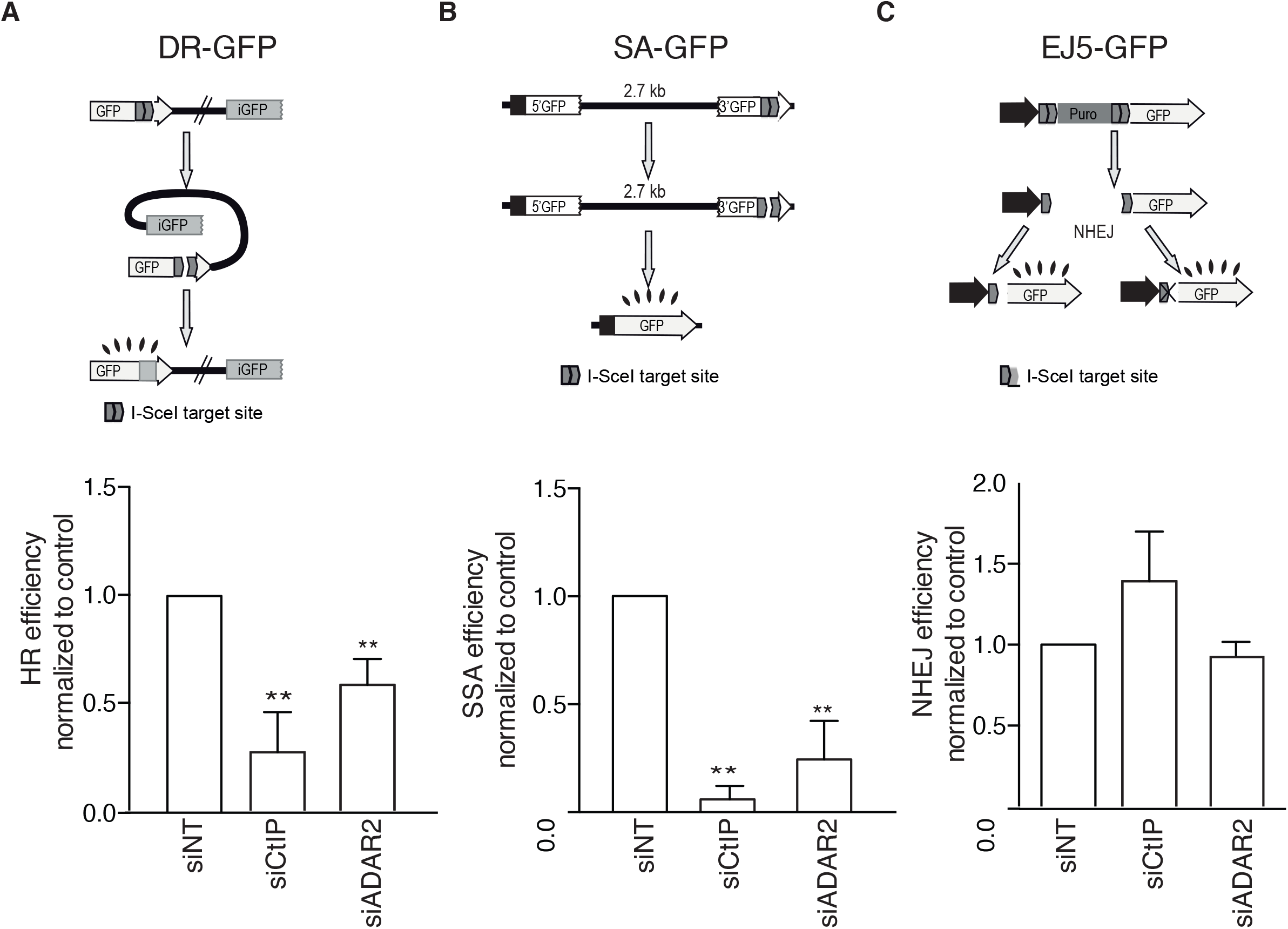
ADAR2 depletion affects homologous recombination. **(A)** Effect of ADAR2 depletion in the DR-GFP reporter. A scheme of the reporter is shown on the top. Induction of a DSB using I-SceI meganuclease render GFP positive cells when the donor repeat (iGFP) is used in a gene conversion event. The efficiency of classical recombination was calculated as the percentage of GFP positive cells in response to I-SceI expression upon down-regulation of the indicated genes and normalized with the control. The average and standard deviation of at least three independent experiments are shown. A Student’s *t*-test comparing cells depleted with a siRNA against ADAR2 with control siNT (Non-target siRNA) was performed and one, two or three asterisk(s) denotes statistical significance at p<0.05, p<0.01 or p<0.001, respectively. **(B)** Same as **A** but using the Single Strand Annealing (SSA) reporter SA-GFP (top). In this case, the induction of a DSB located between two repeats in direct orientation will render GFP positive cells only when intramolecular SSA takes place. **(C)** Same as **A** but using the NHEJ reporter EJ5-GFP (Top). In this case, two I-SceI-induced DSBs could be repaired by conservative or mutagenic NHEJ granting the accumulation of functional GFP.

### DNA resection requires ADAR2 editing activity

Due to its effect in recombination, we hypothesized that ADAR2 might have a role in the common, early steps of the homology-dependent repair pathways, namely in DNA end resection. To test this idea, we first studied RPA foci formation upon ionizing radiation in ADAR2-depleted cells. RPA is a ssDNA binding complex that accumulates at DNA breaks as a direct consequence of DNA end resection^4,5,39^. Thus, the percentage of RPA foci positive cells is the standard readout of resection in mammalian cells. Depletion of ADAR2 in U2OS cells caused a significant defect in resection, though less pronounced than that observed with the downregulation of the key resection factor CtIP (Figure 4A; Supplementary Figure S3A). For representative images of the experiment see Supplementary Figure S3A. The same resection impairment was also observed upon depletion of ADAR2 in HeLa cells (Supplementary Figure S3B). Strikingly, similar results were observed upon depletion of ADAR1, but not the catalytically-dead member of the ADAR family, ADAR3, thus suggesting that active RNA editing is required for DNA end resection (Figure 4A and Supplementary Figure S3A).

**Figure 4.**
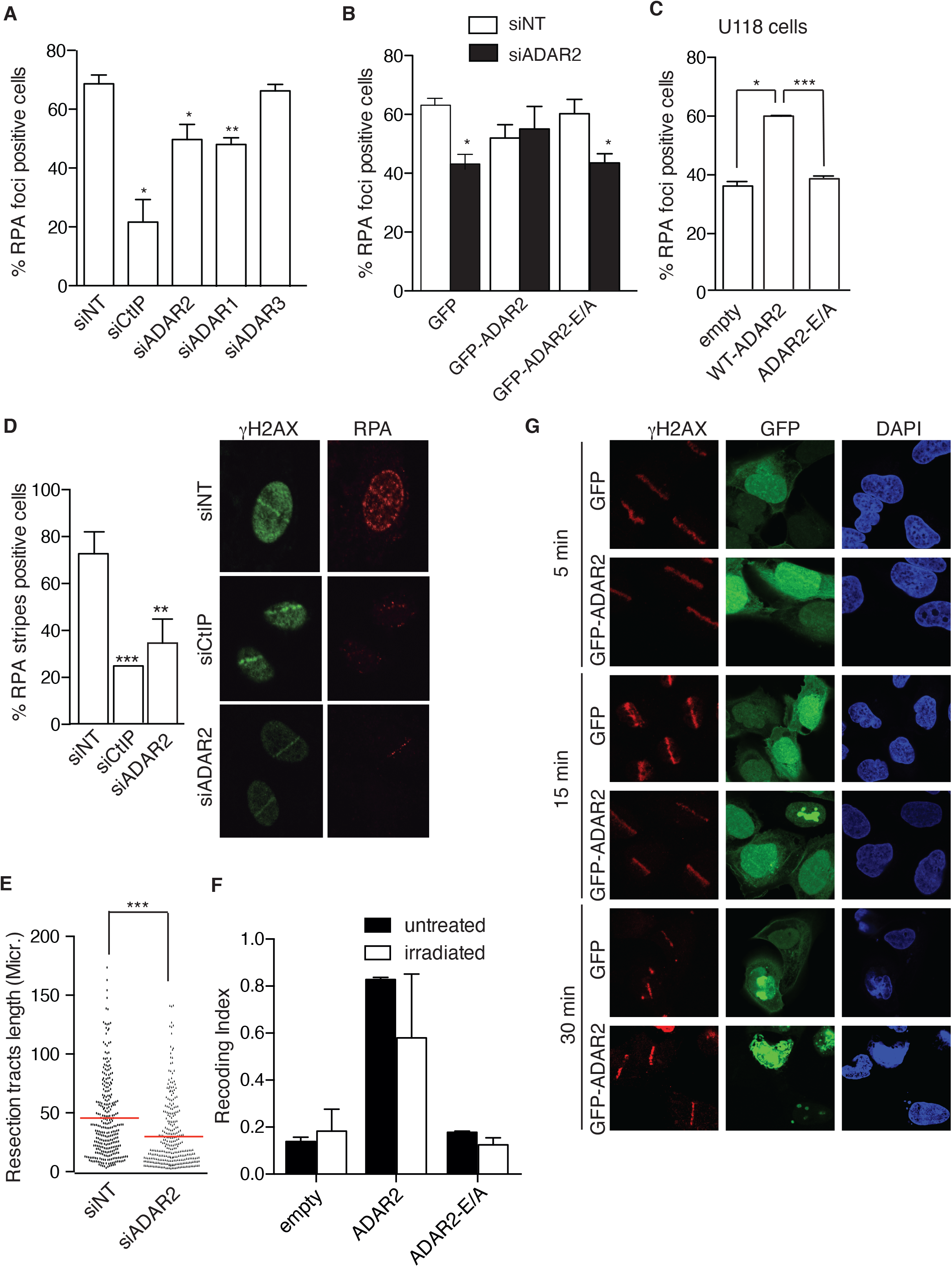
ADAR depletion impairs in DNA resection. **(A)** DNA resection proficiency after 10Gy of irradiation in U2OS cells measured as the percentage of RPA foci-positive cells in cells transfected either with siRNAs against ADAR1, ADAR2, ADAR3, CtIP or with control siNT. The average and SD of three independent experiments are shown. Significance was determined by Student’s *t*-test comparing each condition to siNT cells. *, *P* < 0.05; **, *P* < 0.01; ***, *P* < 0.001. Representative images of the experiments are shown on supplementary figure S3A. **(B)** DNA resection proficiency measured as the percentage of RPA foci-positive cells in U2OS cells expressing either GFP-ADAR2 wild type or a catalytically dead version of the protein (ADAR2 E/A) transfected either with a siRNA against the 3’UTR of ADAR2 (black boxes) or a control siNT-UTR (white boxes). Other details as in **A**. **(C)** RPA foci formation upon in U118 cells treated with 10 Gy of radiation in cells expressing either GFP, GFP-ADAR2 wild type or a catalytically dead version of the protein. Other details as in **A**. **(D)** DNA resection proficiency was measured as RPA stripes–positive cells upon laser micro-irradiation in cells transfected either an siRNA against ADAR2, CtIP or with control siNT. The average and SD of three independent experiments are shown. Representative images of the experiments are shown on the right. Other details as in **A**. **(E)** Resection length measured with the SMART assay using DNA fibres extracted from U2OS downregulated for ADAR2. A non-target siRNA (siNT) was used as control. One out of three representative experiment with similar results is shown. Other details as in (**A**). **(F**) RNA sequencing of U118 cells complemented with a plasmid bearing wild type ADAR2, catalytically dead ADAR2 E/A or the empty vectors in untreated conditions (black bars) or upon exposure to 10 Gy of ionizing radiation (white bars) was used to analyse the changes in RNA sequence of codons known to be edited by ADAR2. The Recoding Editing Index (REI) was reported as percentage. **(G**) U2OS cells bearing GFP-ADAR2 or GFP, as a control, were micro-irradiated using a laser as described in the methods section. Cells were fixed at the indicated time points and the presence of γH2AX (red) or ADAR2 (green) at lasers stripes was analysed. Representative images are shown.

To confirm that the observed phenotype in DNA resection was truly due to the reduction of ADAR2 levels and not to an indirect off-target effect, we studied RPA foci formation in cells bearing siRNA-resistant, GFP-tagged variants of ADAR2. Indeed, the resection impairment caused by depletion of ADAR2 was rescued by wild-type GFP-ADAR2 (Figure 4B and Supplementary Figure S3C). Importantly, this rescue was not observed with the expression of a catalytically-dead version of the protein (GFP-ADAR2E-A) (Figure 4B), arguing that ADAR2 deaminase activity was required for processive resection. Moreover, we could reproduce the resection defect in U118 cells^26^, a glioblastoma cell line that is defective for ADAR2 expression, when compared with the same cells complemented with a wild-type copy of the gene, but not a catalytically-dead mutant (Figure 4C).

To validate these observations, we analyzed recruitment of RPA to DSBs by other means. U2OS cells depleted for ADAR2, or CtIP as a control, were laser micro-irradiated and immunostained with antibodies against RPA and γH2AX to identify the irradiated areas. The percentage of γH2AX-positive stripes that were co-stained by RPA was determined (Figure 4D). In agreement with our previous results, depletion of ADAR2, or CtIP, significantly diminished the presence of RPA at the irradiated areas. Finally, in order to analyse in more detail the resection defect related to ADAR2 depletion and to investigate whether only resection initiation was impaired or if resection processivity was also compromised, we used SMART, a high resolution technique that measures resection in individual DNA fibres^31,40^. As seen in Figure 4E, the length of ssDNA fibres formed during the resection process was reduced upon ADAR2 depletion. Again, similar results were observed upon ADAR1 downregulation (Supplementary Figure S3D), reinforcing the connection between A-to-I editing and DNA end resection.

Altogether, these results confirm that RNA editing by ADAR proteins facilitates DNA end resection at DSBs.

### The role of ADAR2 on resection does not rely on the recoding of mRNAs that encode resection factors

Next, we wondered how this RNA editing activity might be needed for DNA end processing. We studied the recruitment DSB repair factors, such as 53BP1, BRCA1 and CtIP, to DNA damage foci shortly after DNA damage induction upon ADAR2 depletion. Notably, neither of them was affected in cells exposed to ionizing radiation or laser micro-irradiation (Supplementary Figure S4A-C). Then, we wondered if ADAR2 was specifically editing mRNAs that code for resection factors. To analyze this possibility, we exposed the ADAR2-defective cell line U118 to ionizing radiation or mock treatment, isolated total RNA, and sequenced it. U118 cells complemented with either wild-type ADAR2 or a catalytically-dead mutant were also processed in parallel. The levels of ADAR2 mRNA in the different samples are shown in Supplementary Figure S4D. As expected, U118 cells complemented with wild-type ADAR2 showed a higher efficiency in editing the coding codons of known ADAR2 targets, expressed as the weighted average over all known recoding sites, known as the Recoding Editing Index (REI)^35^. Instead, non-complemented U118 cells showed little recoding editing, regardless of whether or not exposed to ionizing radiation (Figure 4F). Equally expected, the expression of a catalytically-dead enzyme also showed almost no recoding of mRNAs (Figure 4F), despite the fact that such variant was expressed almost 3 times more than the wild-type ADAR2 (Supplementary Figure S4D). Thus, only the expression of catalytically-active enzyme led to the expected ADAR2-dependent recoding due to editing of specific codons (Figure 4F). Interestingly, although some specific coding codons were edited more efficiently upon irradiation than in mock treated cells, we observed a general decrease in the recoding editing efficiency of known ADAR2 substrates upon exposure to DNA damage (Figure 4F and Supplementary Table S1). Moreover, we could not observe any change in editing of mRNA from genes that code for recombination or resection factors either in cells exposed to DNA damage or in undamaged cells (Supplementary Table S1). Thus, we conclude that the role of ADAR2 in resection and recombination does not rely on changes in the sequence or expression of specific mRNAs of *bona fide* DNA repair factors. To integrate our data, in which we observed a general ADAR2-mediated increase in RNA editing in the reporter (Figure 1) accompanied by a general reduction of its activity on known targets (Figure 4F), we hypothesized that upon DNA damage ADAR2 is mobilized to induce RNA editing at new sites. Such re-distribution could mean that at least a fraction of the protein would localize at sites of DNA DSBs. To test this idea, we analyzed the presence of GFP-tagged ADAR2 at damaged chromatin at different time points by laser micro-irradiation (Figure 4G). In agreement with our idea, a fraction of GFP-ADAR2 was readily recruited to sites of DNA damage as early as 5 minutes after laser micro-irradiation, something not observed in cells expressing only GFP as a control. In fact, 50% of cells showed colocalization between γH2AX and ADAR2 stripes. Interestingly, such recruitment seemed transient, as it was no longer observed 30 minutes after irradiation. This recruitment pattern agrees with the idea that ADAR2, or at least a fraction of the protein, changes its substrates and is channeled towards RNAs at the sites of broken chromatin to play a role in the early steps of DSB repair.

### ADAR2 facilitates resection over DNA:RNA hybrids

We hypothesized that the ADAR2-depletion related phenotype in DNA resection could be caused by a direct effect of ADAR2 on an RNA molecule located in the vicinity of the break that would act as a physical barrier for DNA end processing. The presence of DNA:RNA hybrids close to DSBs has been documented from yeast to mammals, both as pre-existing R-loops, R-loops formed as a consequence of breaks in transcribed regions or as DNA:RNA hybrids resulting from *de novo* transcription of resected DNA ends^9,13,16,41–43^. The actual effect of such DNA:RNA hybrids in resection is controversial, with both pro- and anti-resection effects described^9,16,44^. Importantly, ADAR2 has been proposed to recognize and edit DNA:RNA hybrids *in vitro*^45^. In order to define whether ADAR2 involvement in DNA end resection depended on the presence of DNA:RNA hybrids, we repeated the resection assay in the presence or absence of ectopically overexpressed RNaseH1, an enzyme that degrades the RNA moiety of such structures^46^. ADAR2 depletion and RNaseH1 overexpression efficiency are documented in Supplementary Figure S5A. Strikingly, the overexpression of RNaseH1 in U2OS reverted ADAR2 resection phenotype as measured by RPA foci accumulation (Figure 5A and Supplementary Figure S5B). Indeed, the mere overexpression of RNaseH1 facilitated RPA foci formation even in cells transfected with a control siRNA, arguing that DNA:RNA hybrids act generally as physical barriers for the resection process, and that ADAR2 helps overcome such roadblocks. To confirm this finding, we studied the recruitment of RPA to damaged chromatin in HeLa cells depleted of ADAR2 and overexpressing RNaseH1. Again, such overexpression rescued the resection phenotype of ADAR2 depletion (Figure 5B). The same was observed, albeit only partially, when laser micro-irradiation experiments were performed (Figure 5C). In none of those cases, the increase in RPA-positive cells was due to changes in cell cycle profile when the RNaseH1 was overexpressed (Supplementary Figure S5C). Then, to assess whether ADAR2 helped remove DNA:RNA hybrids, we analyzed the accumulation of such structures by immunofluorescence using the DNA:RNA hybrid-specific antibody S9.6^47^. As shown in Figure 5D, depletion of ADAR2 increases the nuclear signal with that antibody. To rule out the contribution of other nucleic acid structures to the increase in S9.6 signal, we overexpressed RNaseH1 and observed a significant reduction in the staining (Figure 5D). Of relevance, ADAR2 has been previously found to be a part of the so-called “DNA:RNA hybrid interactome”^48^. Therefore, we decided to test if ADAR2 could interact directly with DNA:RNA hybrids. Indeed, ADAR2 was specifically immunoprecipitated using the S9.6 antibody, in a similar fashion as Senataxin (SETX), a helicase described to dissolve such hybrids and that has been shown to be recruited to DSBs at transcribed regions^13^ (Figure 5E). This immunoprecipitation was specific, as did not occur when a control antibody was used (Figure 5E).

**Figure 5.**
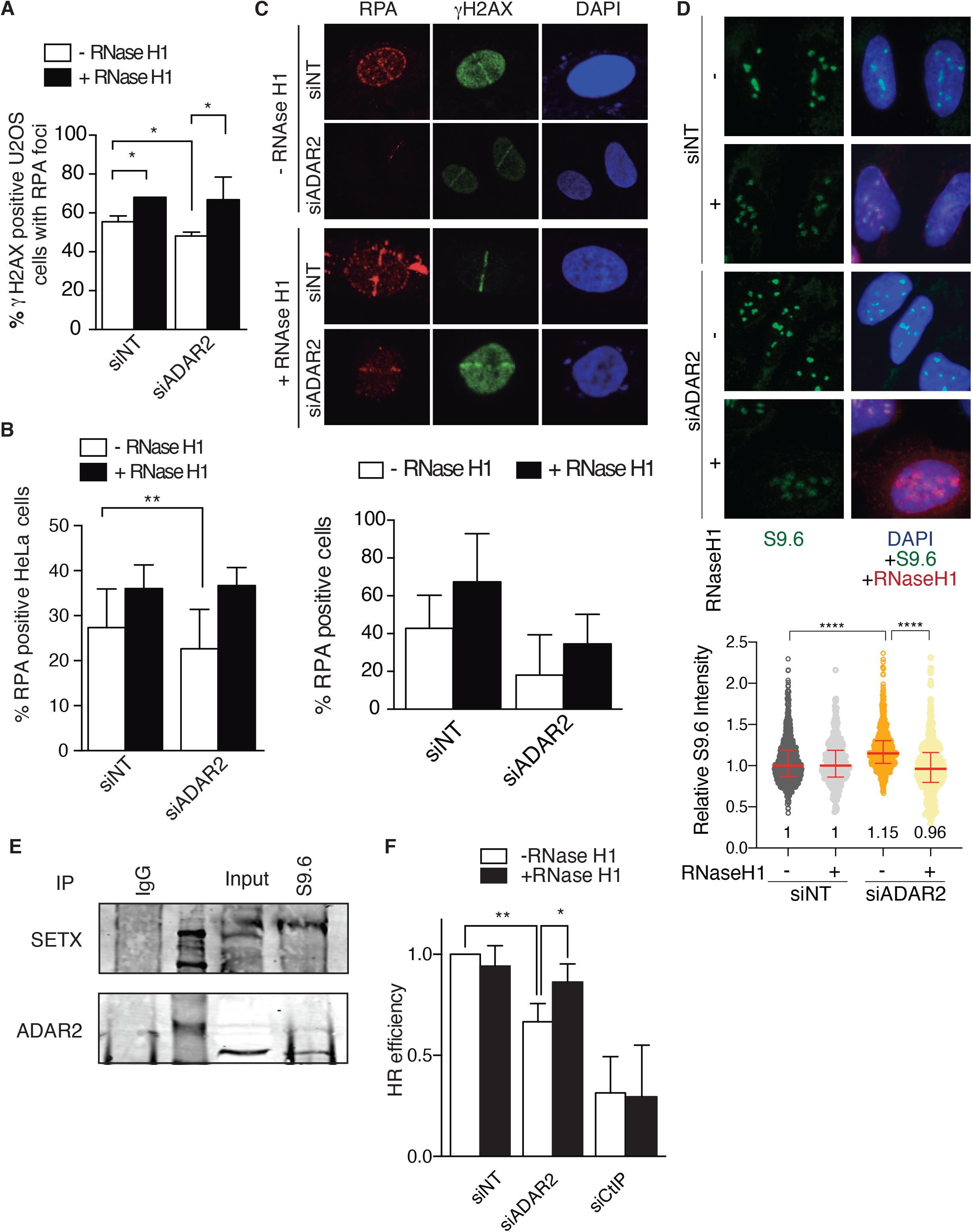
The connection of DNA resection defect with R-loop increase. **(A)** DNA resection proficiency measured as the percentage of RPA foci-positive cells after 1h of 10 Gy of irradiation in cells U2OS transfected either with a siRNA against ADAR2 or with control siNT and transfected either with RNaseH1 overexpression plasmid (black) or with the control empty plasmid (white). The plot shows the percentage of cells positive for RPA foci and the average and standard deviation of at least four independent experiments. For each replicate, at least 200 cells were measured. Other details as figure 4A. **(B)** Same as **A**, but in HeLa cells. **(C)** HeLa cells were transfected with the indicated siRNAs and plasmids and micro-irradiated with a laser to induce DNA damage. Representative images are shown on top. The percentage of cells positive for RPA recruitment to DSB are plotted below the images. The graph shows the average and standard deviation of at least four independent experiments. At least 20 cells per replica were studied and the number of stripes were analysed using FIJI software. (**D**) Accumulation of RNA-DNA hybrids in ADAR2-depleted cells. U2OS cells transfected with siNT and siADAR2 and bearing the pcDNA3-RNaseH1 or pCDNA3 empty vector were immunostained to detect DNA:RNA hybrids using the S9.6 antibody. Relative S9.6 signal intensity per nucleus in U2OS cells with or without overexpression of RNase H1 was calculated (bottom). The median with interquartile range obtained from three independent experiments for each population is shown. Statistical significance was calculated using a Mann–Whitney U-test, ****P < 0.0001. One significative experiment out of three is shown. (**E**) Protein samples from U2OS cells were immunoprecipitated using the anti-DNA:RNA hybrid S9.6 antibody or a non-related IgG as a control. Inputs and immunoprecipitates were resolved in SDS-PAGE and blotted for ADAR2 and Senataxin, as indicated. A representative western blot, out of three independent replicas, is shown. (**F**) Effect of RNaseH1 overexpression in the ADAR2-mediated impairment of recombination. U2OS cells bearing the DR-GFP reporter were transfected with either a siRNA against ADAR2 or with control siNT and either with RNAseH1 overexpression plasmid (white) or with the control empty plasmid (black). The efficiency of classical recombination was calculated as the percentage of GFP positive cells in response to I-SceI expression upon down-regulation of the indicated genes and normalized with the control. The average and standard deviation of three independent experiments are shown. Other details as in figure 3A.

Taken together, our results suggest ADAR2 effect in resection was caused by DNA:RNA hybrid accumulation. A prediction of this model is that the decrease in homologous recombination caused by ADAR2 downregulation should be suppressed by RNaseH1 overexpression. In fact, recombination was almost completely restored in ADAR2-depleted cells when such enzyme was ectopically expressed (Figure 5F). Such effect was not due to a general increase of recombination mediated by RNaseH1 overexpression, as it was not observed either in CtIP depleted or control cells.

### Increase of DNA:RNA hybrids generally impairs DNA end resection

Due to our observation that DNA:RNA hybrids hampered resection in the absence of ADAR proteins, we wondered if other enzymes involved in the removal of hybrids also affected DNA end resection, therefore this represented a more general phenomenon. We decided to test SETX, due to its aforementioned relationship with DSBs. So, we checked if its loss of also affected RPA foci formation. Indeed, depletion of Senataxin also produced a DNA resection defect measured by RPA foci accumulation (Figure 6A) and reduced the length of resected DNA (Figure 6B). As seen for ADAR2 depletion, SETX downregulation also led to an increased burden of spontaneous DNA damage, measured as BRCA1 foci in unchallenged cells (Figure 6C).

**Figure 6.**
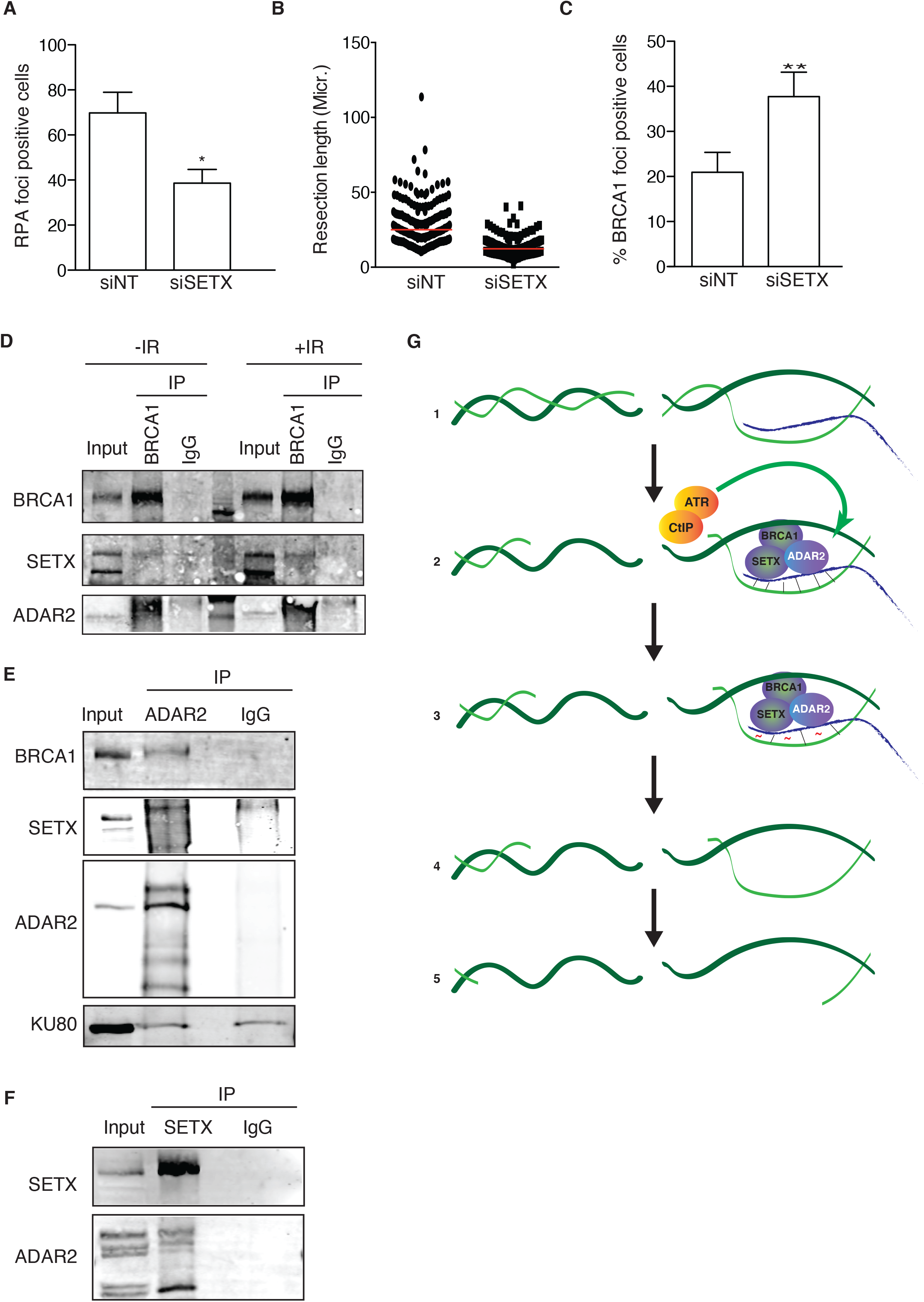
DNA:RNA hybrid stabilization impairs resection. **(A)** Percentage of RPA foci–positive cells in cells transfected either with a siRNA against SETX or with control siNT. Other details as figure 4A. (**B**) DNA resection proficiency measured as the length of resected DNA with SMART in cells transfected either with a siRNA against SETX or with control siNT. Other details as in figure 4E. **(C)** Percentage of BRCA1 foci–positive cells in cells transfected either with a siRNA against SETX or with control siNT. Other details as in figure 1C. **(D)** Protein samples from U2OS cells were immunoprecipitated using an anti-BRCA1 antibody or a non-related IgG as a control, in cells irradiated (left) or not (right). Inputs and immunoprecipitates were resolved in SDS-PAGE and blotted for BRCA1, ADAR2 and Senataxin, as indicated. A representative western blot, out of four independent replicas, is shown. **(E)** Protein samples from U2OS cells were immunoprecipitated using an anti-ADAR2 antibody or a non-related IgG as a control, in non-irradiated cells. Inputs an immunoprecipitates were resolved in SDS-PAGE and blotted for BRCA1, ADAR2, SETX and Ku80 as indicated. A representative western blot, out of three independent experiments, is shown. **(F)** Protein samples from U2OS cells were immunoprecipitated using an anti-Senataxin antibody or a non-related IgG as a control, in non-irradiated cells. Inputs an immunoprecipitates were resolved in SDS-PAGE and blotted for Senataxin and ADAR2, as indicated. A representative western blot, out of three, is shown. **(G)** A schematic representation of how ADAR2 might help resection. (1), DNA:RNA hybrids might appear close to DSBs, either because they were already formed there or specifically formed upon DNA damage. Those hybrids will block resection progression. (2), The formation of some ssDNA by CtIP will activate the ATR branch of the checkpoint, that in turn will stimulate the activity of ADAR2 at DNA:RNA hybrids, including those close to DNA breaks. (3), ADAR2 activity will create mismatches in the DNA:RNA pairing (red tilde), facilitating the dissolution of those structures by SETX-BRCA1. (4-5), Once the DNA:RNA hybrids are eliminated, resection can proceed unimpeded.

### ADAR2 physically interacts with Senataxin and BRCA1

Our data suggested a possible role of ADAR2 in R-loop resolution, which would be key for facilitating DNA end resection and HR. Hence, we wondered if ADAR2 might interact with known players in the homeostasis of R-loops that are also connected with DSB repair. Many different enzymes have been associated with the removal of R-loops. Among them, we decided to focus on the BRCA1-SETX complex. These two proteins have been shown to cooperate in the elimination of R-loops in the 3’ end of many transcribed genes^49^. Moreover, both proteins play roles in DNA end resection, as it has been previously established for BRCA1^40^ and in this study for Senataxin (Figure 6A-C). Thus, we tested whether ADAR2 might physically interact with BRCA1 and Senataxin. Using antibodies against BRCA1, we confirmed the previously shown interaction of this protein with Senataxin^49^ and also established an interaction with ADAR2 (Figure 6D). These interactions were not stimulated by the presence of DNA damage (Figure 6D). Also, by co-immunoprecipitations with different antibodies we could observe the reciprocal interaction with both BRCA1 and SETX (Figure 6E). The possibility that ADAR2 immunoprecipitation could be bringing down any proteins accumulating at sites of DNA double strand breaks was excluded, since we could not observe an interaction with the DNA-end binding protein Ku80, suggesting that ADAR2 interaction with BRCA1 and SETX was indeed specific. It is worth pointing out that these co-immunoprecipitations were performed in the presence of benzonase, thus they are not bridged by DNA. Moreover, immunoprecipitation using an antibody against SETX also allows the co-precipitation of ADAR2 (Figure 6F). Therefore, we could confirm that these three proteins interact in a DNA damage-independent fashion, most likely to facilitate the removal of DNA:RNA hybrids globally.

## DISCUSSION

mRNA post-transcriptional modifications are now being revealed as potent regulators of cellular metabolism that allow flexibility and adaptability to a changing environment^50^. Such chemical modifications of the mRNA can recode coding sequences, create or destroy splicing sites, affect RNA stability and structure or directly interact with specific readers in order to recruit specific machinery affecting translation and/or mRNA localization. Therefore, those modifications affect virtually every aspect of the biological response of the cells. In fact, it has been recently shown that other base modifications of RNA happen at DNA:RNA hybrids and contribute to genomic stability^51^. Thus, it was reasonable to expect that A-to-I RNA editing would contribute to the global response to DNA damage. Indeed, our results support this notion, uncovering a *bona fide* role of A-to-I deamination by ADAR1 and ADAR2 in the maintenance of genomic integrity upon exposure to genotoxic agents. ADAR2-mediated RNA editing seems, indeed, completely upturned upon treatment with DNA damaging agents. Many of the codons in mRNAs that are usually targeted by ADAR2 decrease their editing, whereas specific codons on natural mRNAs and in other RNA species such as the RNAG reporter increase their editing. Based on these findings, we propose an <u>R</u>NA-<u>e</u>diting DNA <u>da</u>mage <u>r</u>esponse (REDAR) that is essential for DNA repair, and specifically for HR, contributing to the maintenance of genomic integrity.

Specifically, we postulate that upon the triggering of the DNA damage response (DDR), REDAR is activated by ATR and CtIP. Then, ADAR2 is mobilized from its usual targets to new ones, most likely including DNA:RNA hybrids, in order to help with their dissolution. In this regard, ADAR2 action might have two non-mutually-exclusive outcomes; on the one hand it can rapidly and transiently relocate to promote RNA editing at the sites of DSBs, aiding in the removal of DNA:RNA hybrids and facilitating DNA end resection; and on the other hand, it can increase the editing of a small fraction of yet undisclosed mRNAs, whose role in the DDR should be clarified in further studies. Interestingly, it has been previously shown that ADAR action can alter the biochemical properties of some DNA repair enzymes^23^. We envision that the transcriptional inhibition induced by the DDR^52^ will cause a reduction of the normal co-transcriptional RNA editing^53^ and will free ADAR to act on alternative substrates such as the aforementioned DNA:RNA hybrids. We favor the idea that the editing itself would happen on the RNA moiety of the structure, as we could not observe accumulation of mutations compatible with ADAR activity on DNA (AT-to-GC pair changes) in DiVA cells neither in unchallenged cells nor upon induction of DSBs. DiVA cells are modified U2OS cells in which DSBs can be created at specific and well mapped sites by the translocation of the restriction enzyme *Asi*SI from cytoplasm to nucleus^54^. In fact, similar results were obtained in cells depleted for ADAR2, overexpressing this factor or in control cells (data not shown). However, by performing an *in silico* study we observed that cancer cells that overexpress ADAR2 are statistically more likely to accumulate this kind of mutations (Supplementary Figure S6). Thus, it is also possible that the editing occurs in the DNA strand, as suggested *in vitro*^45^, albeit how frequently and whether this happens in the context of DNA repair is still unclear. Editing of the DNA strand would greatly increase the mutagenesis associated with HR, something that might be deleterious for the cells, but has been observed due to the action of C-to-U deaminases in cancer^55^. Interestingly, the accumulation of N6-methyladenosine modification on the RNA at DNA:RNA hybrids has also been shown to ease the dissolution of such structures^51^. As A-to-I editing is negatively influenced by m^6^A modifications^56,57^, the crosstalk between those two RNA post-transcriptional modification during DNA repair and in response to DNA damage will be worth exploring further.

The mechanism by which ADAR2 is relocated in response to DNA damage is still far from clear. On the one hand, a passive model is possible, in which ADAR2 naturally recognizes and binds to DNA:RNA hybrids. It has been shown that DNA damage induces the formation of such structures and, in this scenario, the accumulation of hybrids would sequester ADAR2, reducing its availability to edit its usual targets. Indeed, using GFP-tagged versions of ADAR2 we could observe a re-localization to laser micro-irradiated stripes. However, such accumulation is very transient, and in a completely different timeframe of the changes in recoding we have also shown. So, this recruitment cannot fully explain the changes in editing we observed globally, arguing for the coexistence of both a global, genome-wide effect, and a site-specific role. It is even possible that, in this context, during REDAR, ADAR2 would not act specifically at DNA:RNA hybrids that are close to DSBs, but rather relocated to hybrids spread through the genome as a broad response to DNA damage. In this scenario, the accumulation at laser lines we observed would rely simply on the already known accumulation of hybrids at break sites. Indeed, the SETX-BRCA1 complex has been proposed to be important and recruited mainly to transcription termination sites, regardless of the presence of DNA damage^49^, thus again arguing with for putative role that does not require specific recruitment to DSBs. Alternatively, there could be an active recruitment of ADAR2 to hybrids that sit specifically in the proximity of DSBs. The physical interaction with BRCA1, a well-established, *bona fide* member of the DNA damage response, which is recruited to DSBs^58^ and Senataxin, that is also localized to damaged chromatin^13^, might favor this model. This idea is also supported by the fact that A-to-I editing at new sites is upregulated upon REDAR activation in an ATR dependent manner, arguing for a DNA damage-dependent recruitment of ADAR2 to those new target sites. Interestingly, the ATR checkpoint is mainly active in S and G2, as it requires ssDNA for its activation that is produced by DNA end resection. This would explain why CtIP downregulation also decreases the DNA damage-induced increase of RNA editing in the RNAG system, simply reflecting the role of CtIP in ATR activation upon formation of DSBs.

CtIP is linked to RNA metabolism in multiple ways. It affects RNA editing, as shown here, but also interacts with multiple RNA binding proteins^59^ that in turn are required for proper resection. Moreover, we have recently shown that CtIP controls the splicing of specific factors, in many cases facilitating the accumulation of specific alternative splicing forms upon exposure to DNA damage^60^. Additionally, CtIP not only interacts with BRCA1, which also affects RNA splicing, but also CtIP deficiency has been shown to promote the accumulation of DNA:RNA hybrids at sites of highly expressed genes^14^. Paradoxically, CtIP depletion reduces DNA:RNA hybrid accumulation dependent on *de novo* transcription of dilncRNA (damage-induced long non-coding RNAs) starting at DSBs^42^. Hence, CtIP depletion seems to increase R-loops that are produced as a consequence of previous transcription and appears to decrease *de novo* production of diRNAs (DSB-induced small RNA), thus reducing the DNA:RNA hybrids formed after DNA damage. The data suggest that DNA resection, and specifically CtIP and BRCA1, are in close relationship with the general metabolism of RNAs, a reciprocal connection that is worth to continue exploring.

To integrate all our observations, we present a model in which ADAR2, BRCA1 and SETX facilitate the removal of hybrids genome-wide even in the absence of DNA damage, but in a way that is stimulated by the DDR during REDAR. When exposed to a source of DSBs, the levels of DNA:RNA hybrids and R-loops increase, as previously described by other authors^13–16^, and many of them specifically accumulates at break sites. Although some authors argue that these structures might favor the repair process^44^, other favor a view in which they act as roadblocks for repair (for a review see ^43^). This contradiction might simply stem from a differential effect of DNA:RNA hybrids depending on the timing of repair, i.e. very early events might need them but later they have to be eliminated, or the position of the hybrids, as discussed elsewhere^9,61^. Our data agree with the notion that at least some of them represent roadblocks for the progression of the repair machinery that have to be removed prior to repair (Figure 6G-1). Such blocking effect of R-loops is well established for other DNA transactions such as transcription and replication^43,62^. We propose that, upon irradiation, a CtIP- and ATR-mediated global induction of A-to-I editing at DNA:RNA hybrids facilitate their removal both genome-wide and locally, at sites of DNA breaks, hence permitting the DNA end resection machinery to overcome the physical barrier represented by pre-existing or recently formed hybrids close to the DNA breaks (Figure 6G-2). Mechanistically, we suggest that ADAR2-mediated editing of DNA:RNA hybrids, an activity that has been confirmed *in vitro*^45^, might alter the sequence of the RNA strand creating ribo-Inosine (rI) and the deoxiribo-thymine (dT) mismatches (Figure 6G-3). The appearance of those mismatches will loosen up the interaction between the RNA and the DNA strand, facilitating the unwinding of the structure by the helicase activity of SETX (Figure 6G-3,4), allowing the resection machinery to go through (Figure 6G-4,5). Strikingly, ADAR proteins were first discovered in *Xenopus* as having developmentally regulated dsRNA unwinding activity in oocytes^63^, and later shown to rely on the modification of A-to-I in the RNA substrate, which modified the base pairing properties and facilitated the melting of those RNA:RNA double-stranded structures^64^. Alternatively, rather than physically loosening the interaction between the DNA and RNA, A-to-I modification of the RNA might help the recruitment of proteins involved in R-loop removal or even the recuitment of *bona fide* resection factors. We propose that this alteration of the RNA moiety of the hybrids will happen at all R-loops scattered across the genome, even in the absence of DNA damage. But particularly at DSBs, ADAR2 will contribute to the processivity of resection over DNA:RNA hybrids, regardless if these structures preceded or formed as consequence of the break in its vicinity. Our model proposes a role of ADAR2 in DNA end resection and recombination that acts independently, but not exclusively, of the recoding or regulation of expression of yet-undisclosed mRNAs.

Strikingly, ADAR deficiency, and generally unbalance of the levels of editing of the RNA, has been associated with the development of many different tumors^24,25,35,65^. Specific editing events have already been associated with glioblastoma^22,66^. However, in the light of this new role for ADAR2 in DNA repair and the DDR, it might be important to extend the connection of this protein with cancer and establish if it is linked to a particular genomic instability signature.

## ACKNOWLEDGEMENTS

This work was funded by a R+D+I grant from the Spanish Ministry of Economy and Competitiveness (SAF2016-74855-P to P.H. and BFU2016-75058-P to A.A.), the R+D+I grant PID2019-104195G from the Spanish Ministry of Science and Innovation-Agencia Estatal de Investigación/10.13039/501100011033 (P.H.), the European Research Council (ERC2014 AdG669898 TARLOOP to A.A.), the Associazione Italiana Ricerca sul Cancro-AIRC (AIRC IG 22080 to A.G.), The Swedish Research Council (grants 2015-04553 to N.V.), The Swedish Cancer Society (grant CAN 2016/460 to N.V.) and the European Union (FEDER). AR is, and FM-N and RP-C were, funded with FPU fellowships from the Spanish Ministry of Education and S.S. by a Juan de la Cierva contract from the Spanish Ministry of Economy and Competitiveness. J.D.P. and M.E-C. were supported by the Department of Molecular Biosciences, the Wenner-Gren Institute at Stockholm University. J.D-P. and R.P-C. were recipients of short-term EMBO fellowships (STF-7513-2018 and STF-7764-2018). The mutational results shown here are in whole or part based upon data generated by the TCGA Research Network: https://www.cancer.gov/tcga. All the bioinformatic analyses were performed using custom scripts that were run on the High Perfomance Computing cluster provided by the Centro Informático Científico de Andalucía (CICA). CABIMER is supported by the regional government of Andalucia (Junta de Andalucía).

## DATA AVAILABILITY

All relevant data are included in the manuscript. Raw data will be provided upon request.

