## Supplementary figures and images for "ADAR2-mediated RNA editing of DNA:RNA hybrids is required for DNA double strand break repair"

### Supplementary Figure 1

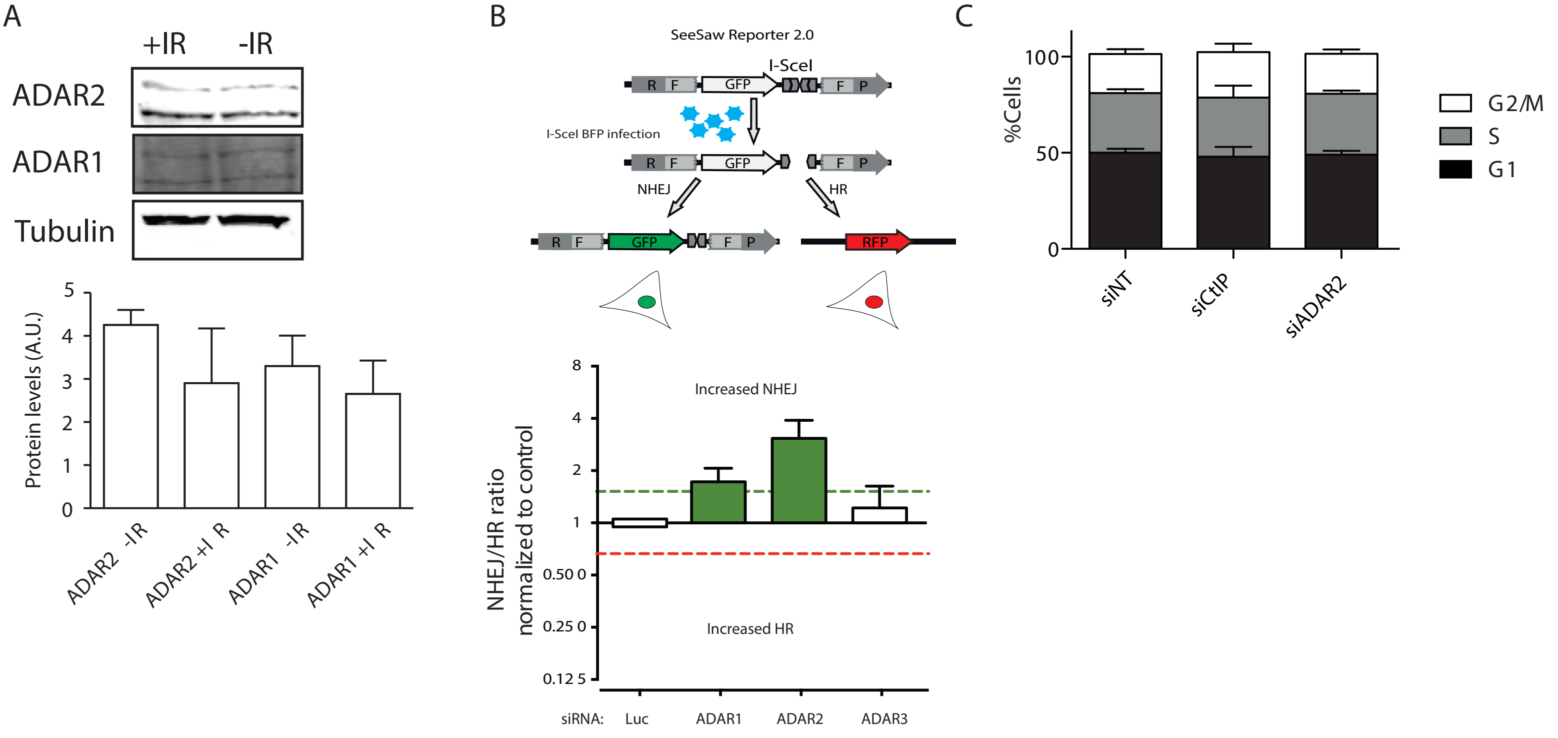


Jimeno et.al. Figure S1

### Supplementary Figure 2

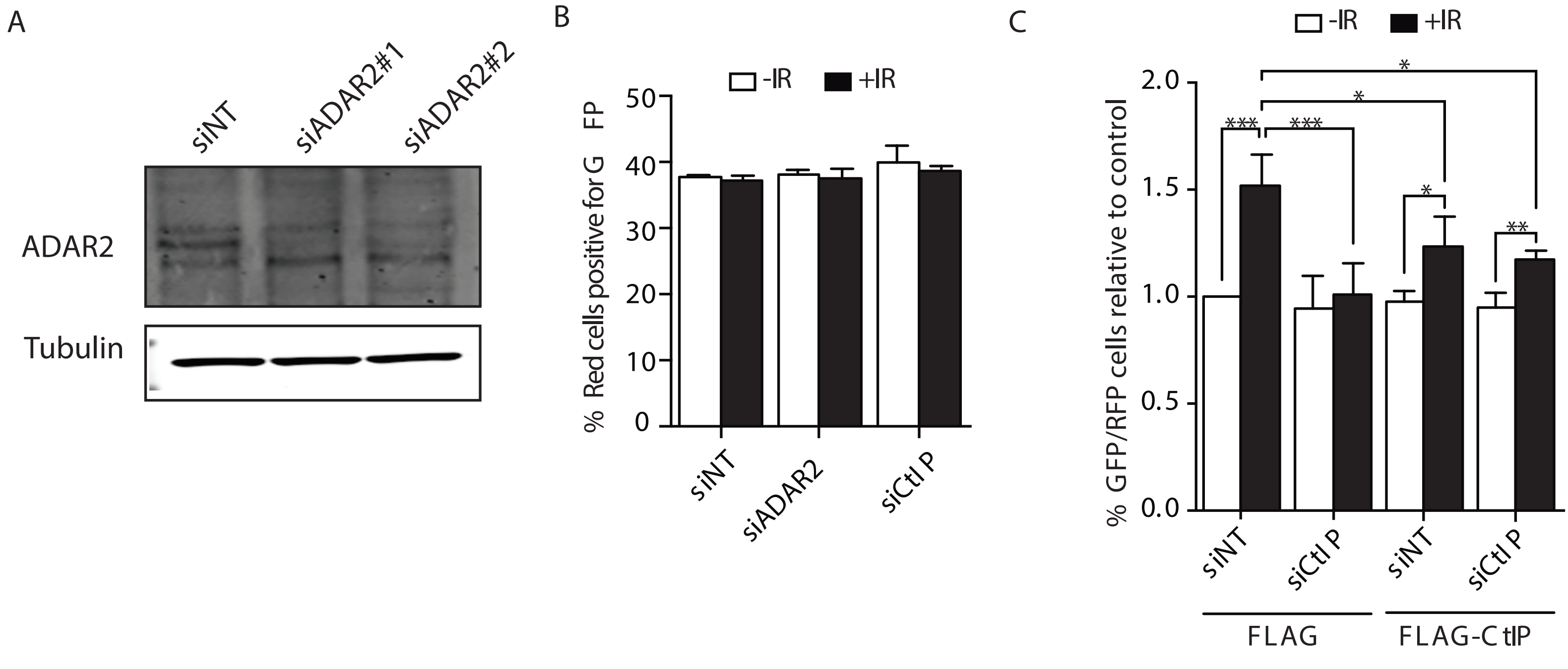


Jimeno et.al. Figure S2

### Supplementary Figure 3

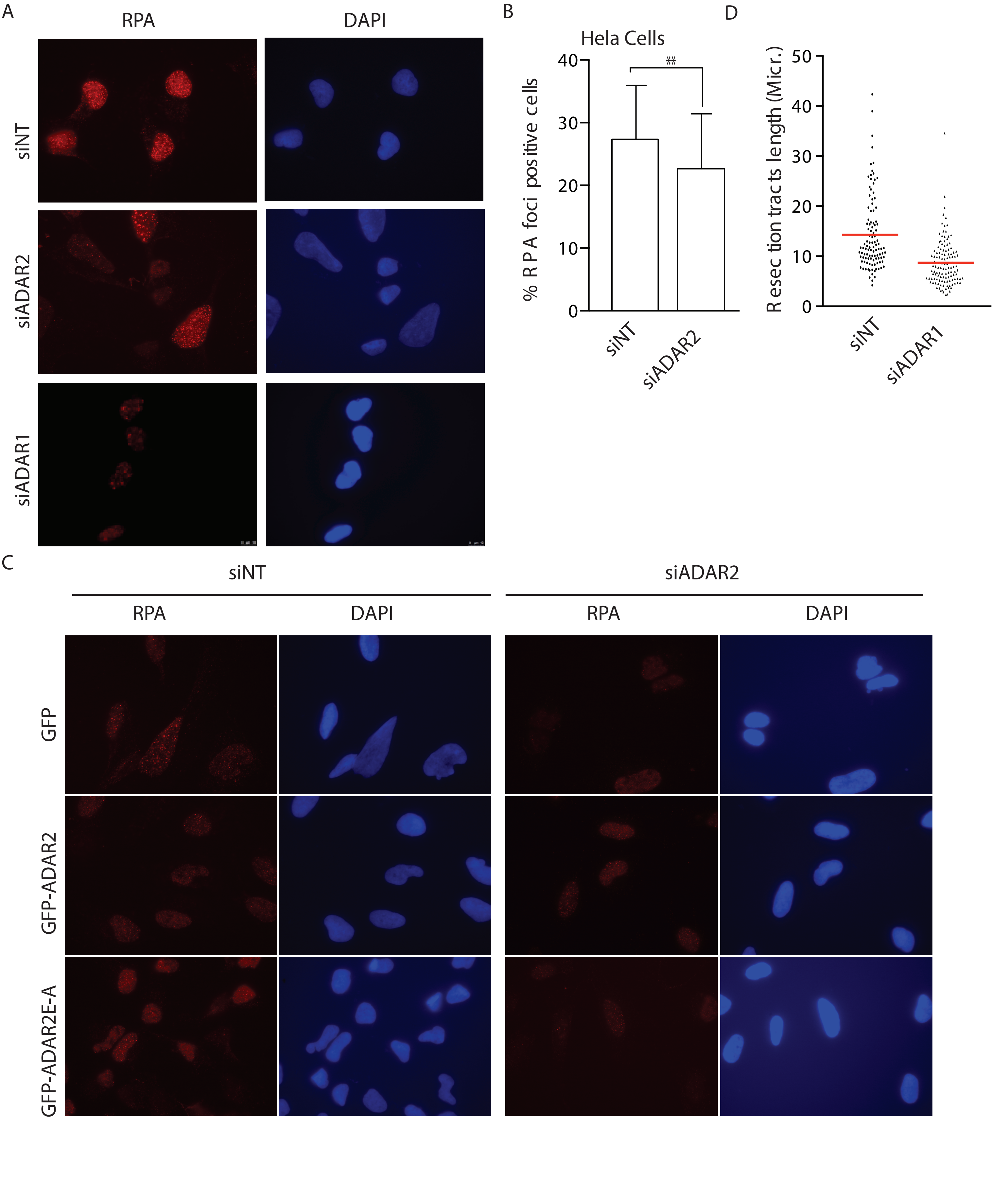


Jimeno et.al. Figure S3

### Supplementary Figure 4

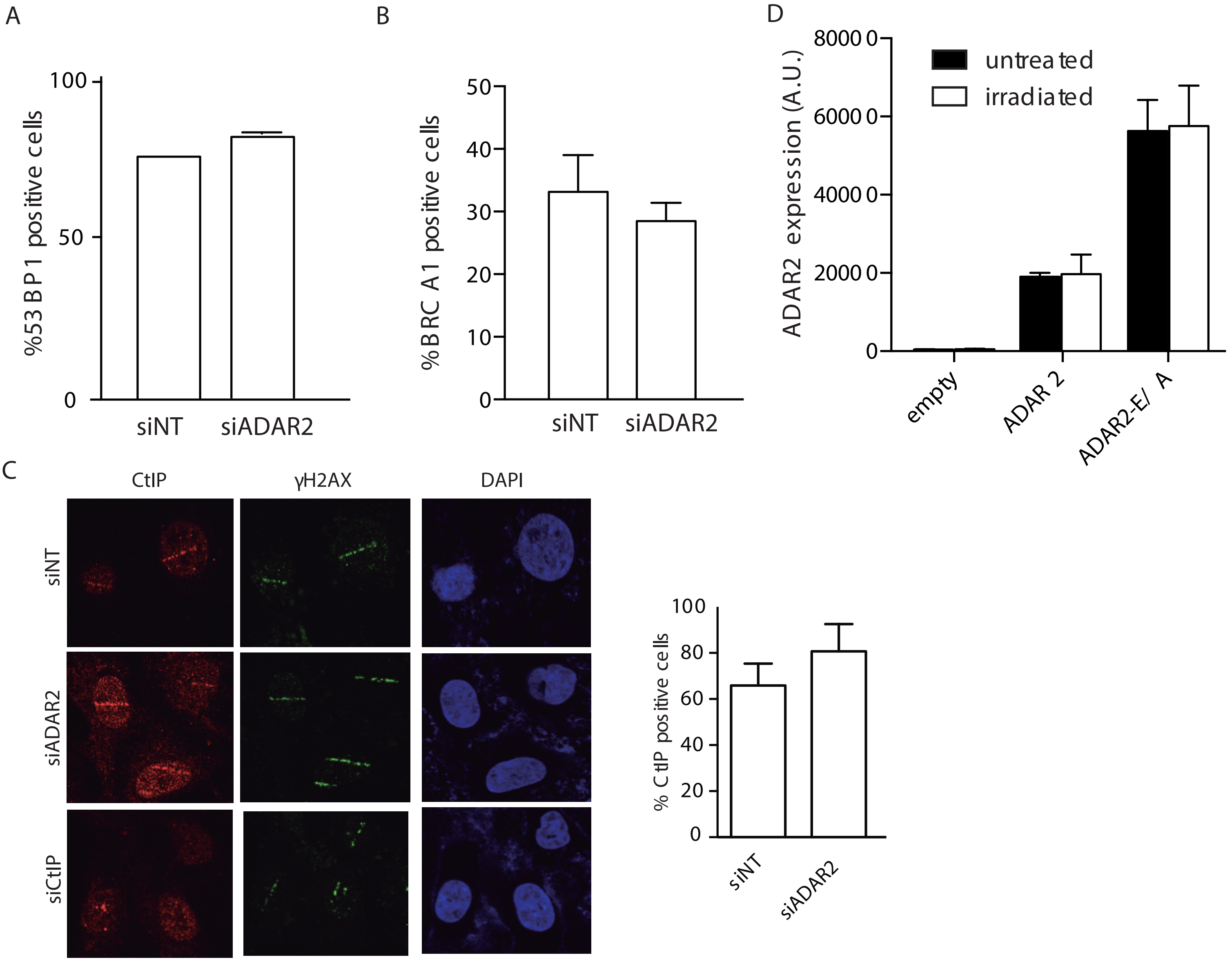


Jimeno et.al. Figure S4

### Supplementary Figure 5

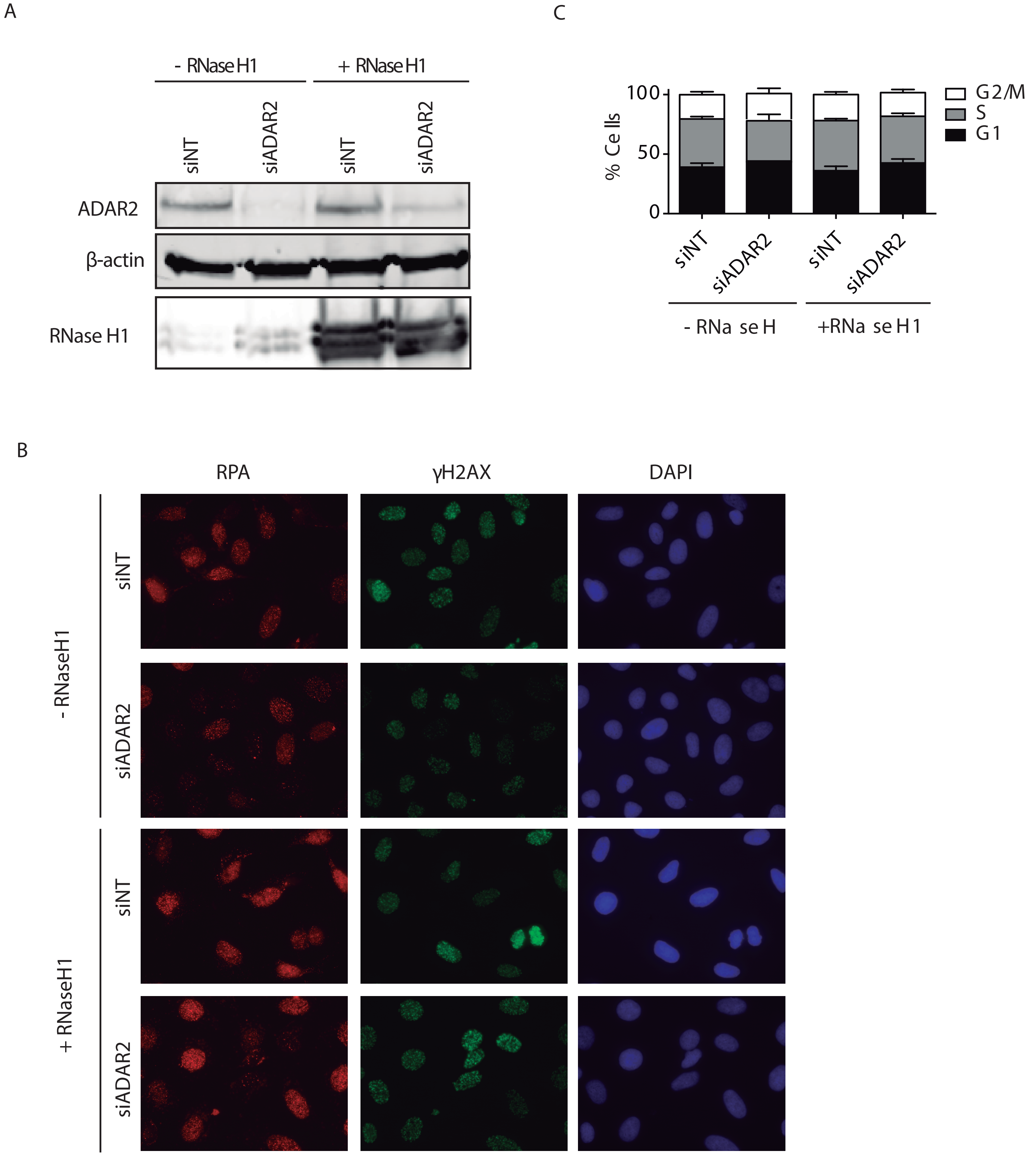


Jimeno et.al. Figure S5

### Supplementary Figure 6

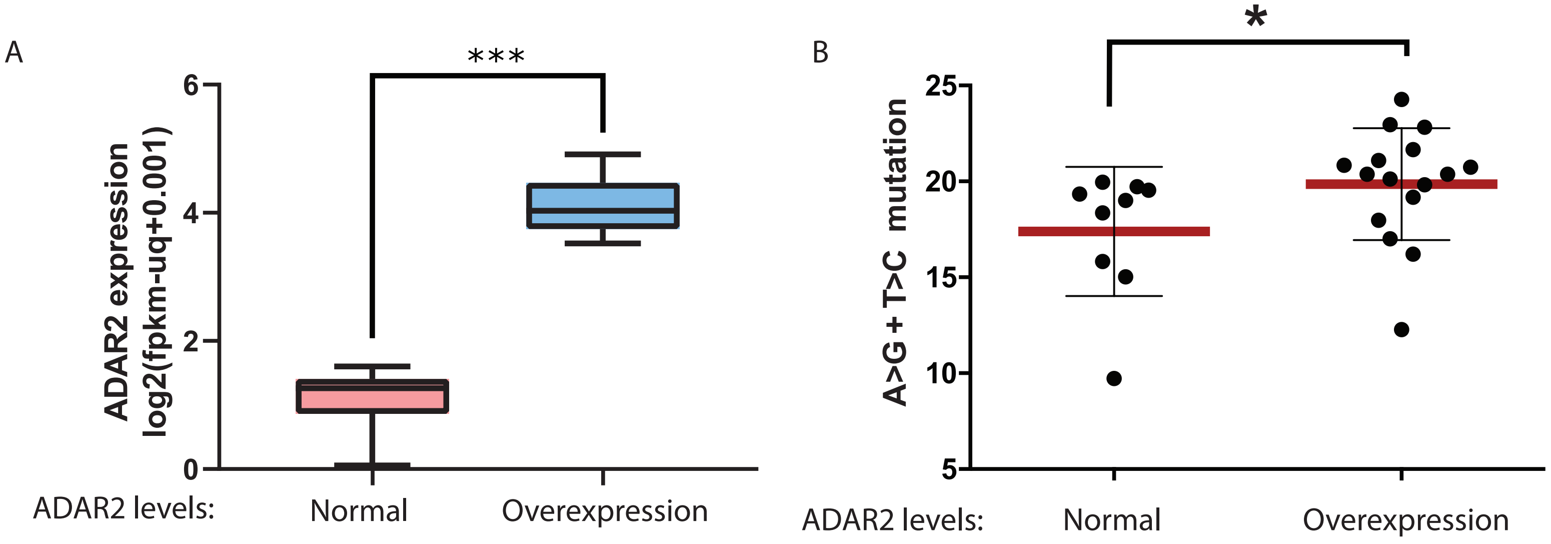


Jimeno et.al. Figure S6
