## Supplementary Information for "ADAR2-mediated RNA editing of DNA:RNA hybrids is required for DNA double strand break repair"

**SUPLEMENTAL MATERIAL:**

**EXPERIMENTAL PROCEDURES:**

**Table S2. siRNAs used in this study.**

| Target gene | Description | Source/Reference/Sequence |
|---|---|---|
| *ADAR2* | SiADAR2-1 (#1) | CCAUUUACUUCUCGAGCAU |
| *ADAR2* | SiADAR2-3(#2) | CGCAGAGUUCCUCACUGUA |
| *ADAR2* | Si ADAR2-3´ÚTR | AAAGCACAGUCUAUGGAACGCUAAT |
| *CtIP* | SiCtIP | GCUAAAACAGGAACGAAUC |
| *SETX* | SiSETX | GCCAGAUCGUAUACAAUUA |
| *NT* | *Non-target* | Sigma, SHC016 |
| *ADAR3* | *siADAR3* | CCUUAAAAUAUGAAUUUACAUGUTA |
| *ADAR1* | *siADAR1* | CAGAUAAUCAUGACUUAGCAAGAAT |

**Table S3. Primary antibodies used in this study.** WB, western blotting. IF, immunofluorescence. SMART, Single Molecule Analysis of Resection Tracks. IP, immunoprecipitation.

| Primary antibody | Supplier | Reference | Application | Concentration |
|---|---|---|---|---|
| **BrdU (mouse)** | Sigma Aldrich | RPN202 | SMART | 1:100 |
| **GFP (rabbit)** | Santa Cruz | sc-8334 | IF, WB | 1:250 |
| **BRCA1 (mouse)** | Santa Cruz | sc-6954 | WB, IF, IP | 1:500 |
| **CtIP (mouse)** | Santa Cruz | Gift from Richard Baer | IF | 1:500 |
| **γH2AX (mouse)** | Cell Signaling | 2577L | IF | 1:250 |
| **ADAR2 (mouse)** | Santa Cruz | sc-73409 | WB, IP | 1:500 |
| **RPA2 (mouse)** | Abcam | ab2175 | IF | 1:500 |
| **53BP1 (rabbit)** | Novus | NB100-304 | IF,WB | 1:500 |
| **SETX (rabbit)** | Bethyl Laboratories | (A301-105A) | WB, IP | 1:1000 |
| **RNAseH1** | Proteintech, | (156061-AP) | WB | 1:1000 |
| **α-tubulin (mouse)** | Sigma | T9026 | WB | 1:50000 |
| **β-actine (rabbit)** | Abcam | Ab8227 | WB | 1:50000 |
| **S9.6 (D5H6) (mouse)** | Covalab | Mab0105-P | IF/IP | |
| **Ku80 (mouse)** | Invitrogen | MA1-23314 | WB | 1:500 |

**Table S4. Secondary antibodies used in this study.** WB, western blotting. IF, immunofluorescence. SMART, Single Molecule Analysis of Resection Tracks.

| Secondary antibody | Supplier | Reference | Application | Concentration |
|---|---|---|---|---|
| **Alexa Fluor 594 goat anti-mouse** | Invitrogen | A11032 | IF, SMART | 1:500 |
| **Alexa Fluor 488 goat anti-rabbit** | Invitrogen | A11034 | IF | 1:500 |
| **Alexa Fluor 594 goat anti-mouse** | Jackson Immunorese | 115-585-003 | IF (laser line) | 1:100 |
| **Donkey anti-rabbit-FITC** | Jackson Immunoresearch | 711-096-152 | IF (laser line) | 1:100 |
| **IRDye 680RD goat anti-mouse IgG (H+L)** | Li-cor | 926-68070 | WB | 1:5000-1:10000 |
| **AIRDye 800RD goat anti-rabbit IgG (H+L** | Li-cor | 926-32211 | WB | 1:5000-1:10000 |

### SUPPLEMENTARY FIGURE LEGENDS

**Supplementary Figure S1. ADAR2 unbalance the repair of DSBs.** (**A**) DNA damage does not induce ADAR1 or ADAR2 accumulation. Protein samples from U2OS cells bearing the RNAG reporter collected 1 h after 10 Gy of ionizing radiation (+IR) or mock treated (-IR) were resolved in SDS-PAGE and blotted with antibodies against ADAR1 and ADAR2 or tubulin as a loading control. The plot shows the media of the quantification of protein levels of two different experiments. (**B**) In the top side, a schematic representation of the SeeSaw reporter (SSR). Infection with a virus bearing the *I-SceI* gene induces the formation of a DSB in the reporter. When repaired by NHEJ, an active *GFP* gene is reformed. However, the break can be also repaired by intra-strand recombination using two copies of truncated RFP. In that case, the cells will express the RFP protein and lose the *GFP* gene. On the bottom side, the effect of the depletion of different ADAR family members in the SSR is shown. Data from (*1*) (**C**) Cell cycle analysis of U2OS cells transfected with the indicated siRNAs measured by FACs. The average and standard deviation of three independent experiments is shown.

**Supplementary Figure S2. DNA damage induced RNA editing.** (**A**) Representative western blot showing siRNA-mediated depletion of ADAR2. U2OS cells were transfected with the indicated siRNAs. 48h later protein samples were obtained, resolved in SDS-PAGE and blotted with the indicated antibodies. (**B**) Effect of ADAR2 and CtIP depletion in the control reporter RNWG. Details as in Figure 1D. (**C)** Effect of CtIP ectopic expression on DNA damage-induced RNA editing. U2OS bearing the RNAG reporter system were transfected with siCtIP or a control siRNA (siNT) and complemented with a FLAG-CtIP construct or a FLAG empty vector. Other details as in Figure 1.

**Supplementary Figure S3. ADAR2 depletion impairs DNA end resection.** (**A)** Representative images of the experiment shown in Figure 4A. (**B**) RPA foci in HeLA cells depleted for ADAR2. Other details as in Figure 4A. (**C)** Representative images of the experiment shown in Figure 4B. (**D**) DNA end resection upon downregulation of ADAR1 using SMART. Other details as in figure 4E.

**Supplementary Figure S4. ADAR2 depletion does not grossly affects the recruitment of pro- or anti- resection factors.** (**A)** U2OS cells transfected with the indicated siRNAs were irradiated (10 Gy). One hour later, cells were prepared for immunostaining using a 53BP1 antibody. The number of 53BP1 foci-positive cells is plotted. Graph represent the average and standard deviation of three independent experiments. (**B**) Same as A, but using a BRCA1 antibody. (**C)** Cells were laser-microirradiated as described in the methods section and stained with an antibody against CtIP. Representative images are shown on the left, and the average and standard deviation of three independent experiments is shown on the right. (**D**) Expression levels of ADAR2 in U118 cells complemented with a plasmid bearing the wildtype gene (ADAR2), a catalytically dead mutant (E/A) or the empty vector. Expression was obtained by RNA sequencing. The average and standard deviation of two independent experiments is shown.

**Supplementary Figure S5. ADAR2 resection upon overexpression of RNase H1.** (**A)** Representative western blot of ADAR 2 depletion and RNAseH1 overexpression. U2OS cells were transfected with pcDNA3-RNaseH1 or pCDNA3 and depleted or not of ADAR2. Protein samples were isolated, resolved in SDS-PAGE and blotted with the indicated antibodies. (**B**) Representative images of the experiment shown in Figure 5A. (**C)** Cell cycle distribution of cells treated as in (A) and measured by FACs analysis. See methods for details.

**Supplementary Figure S6. Mutagenesis associated to ADAR2 overexpression in cancer cell lines.** (**A)** Stratification of tumor samples regarding ADAR2 expression levels in the control group (ADAR2 WT, n=17) and overexpressing group (ADAR2 +, n=24). Expression levels are expressed as log2(fpkm-uq+0.001). (**B**) A to G and T to C somatic mutation levels in tumors from the ADAR2 overexpressing (ADAR2 +, n=24) and ADAR2 control group (ADAR2 WT, n=17). A to G and T to C percentage were used as a measurement for ADAR2 impact over the mutational landscape of ADAR2. A to G and T to C percentage was calculated using the total amount of mutations in each donor set. Statistical significance was calculated with a Student's *t*-test with Welsch correction and is denoted with one ($p<0.05$) or three ($p<0.001$) asterisks.
